# Massive windborne migration of Sahelian insects: Diversity, seasonality, altitude, and direction

**DOI:** 10.1101/2020.02.28.960195

**Authors:** J Florio, L Verú, A Dao, AS Yaro, M Diallo, ZL Sanogo, D Samaké, DL Huestis, O Yossi, E Talamas, L Chomorro, JH Frank, M Biondi, C Morkel, C Bartlett, Y-M Linton, E Strobach, JW Chapman, DR Reynolds, R Faiman, BJ Krajacich, CS Smith, T Lehmann

**Affiliations:** Laboratory of Malaria and Vector Research, NIAID, NIH. Rockville, MD, USA; Malaria Research and Training Center (MRTC) / Faculty of Medicine, Pharmacy and Odonto-stomatology, Bamako, Mali; Systematic Entomology Laboratory - ARS, USDA, Smithsonian Institution, National Museum of Natural History, Washington, DC, USA; Florida Department of Agriculture and Consumer Services, Division of Plant Industry, Gainesville, FL, USA; Entomology and Nematology Department, University of Florida, Gainesville, FL, USA; Department of Life, Health, and Environmental Sciences, University of L’Aquila, Italy; Institute of Applied Entomology, Beverungen, Germany; Department of Entomology and Wildlife Ecology, University of Delaware, Newark DE, USA; Earth System Science Interdisciplinary Center, University of Maryland, College Park, MD, USA; Walter Reed Biosystematics Unit, Smithsonian Institution Museum Support Center, Suitland, MD, USA and Department of Entomology, Smithsonian Institution, National Museum of Natural History, Washington, DC, USA; Centre for Ecology and Conservation, and Environment and Sustainability Inst., University of Exeter, Penryn, Cornwall, UK and College of Plant Protection, Nanjing Agricultural University, Nanjing, P. R. China; Natural Resources Institute, University of Greenwich, Chatham, Kent, ME4 4TB, UK, and Rothamsted Research, Harpenden, Hertfordshire AL5 2JQ, UK; American Museum of Natural History, New York, NY, USA

**Keywords:** aero-ecology, Africa, biodiversity, conservation, dispersal, food security, insect-migration, public health, Sahel, windborne

## Abstract

Knowledge on long-distance migration of insects is especially important for food security, public health, and conservation–issues that are especially significant in Africa. During the wet season, the Sahel nourishes diverse life forms which are soon purged by the long dry season. Windborne migration is a key strategy enabling exploitation of such ephemeral havens. However, our knowledge of these large-scale movements remains sparse due to the virtual invisibility of insects flying at altitude. In this first cross-season investigation (3 years) of the aerial insect fauna over Africa, we sampled crepuscular and nocturnal insects flying 40–290 m above ground in four Sahelian villages in Mali, using sticky nets mounted on tethered helium-filled balloons. Nearly half a million insects were caught, representing at least thirteen insect orders following preliminary sorting of the collections. At least 100 insect families were determined to have been captured at altitude in samples collected on 222 nets, obtained in 125 collections over 96 nights. Control nets (raised momentarily to >40 m during system launch and retrieval) confirmed that the insects were captured at altitude, not near the ground. Thirteen ecologically and phylogenetically diverse species were studied in detail. The flight activity of all species peaked during the wet season every year across localities up to ~100 km apart, and occurred over multiple nights, suggesting regular migrations. Species differed in flight altitude, seasonality, and correlations with aerial temperatures, humidity, and wind speed. All taxa exhibited frequent migrations on southerly winds, accounting for the recolonization of the Sahel from southern source populations. “Return” southward movement at the end of the wet season occurred in most taxa but no selectivity for such winds was detected. Extrapolation of aerial density to estimate the seasonal number of migrants crossing Mali at latitude 14°N suggested numbers in the trillions, even for the modestly abundant taxa. Assuming 2–10 hours of flight, the nightly distances traversed exceed tens and even hundreds of kilometers. Two migration strategies were proposed: “residential Sahelian migration” and “round trip migration”. The unprecedented magnitude and diversity of long-range windborne insect migrations highlight the importance of this life strategy in their impact on Sahelian and neighboring ecosystems.

## INTRODUCTION

Long-distance migration of insects has been recognized as a pivotal influence affecting food security (Glick, 1939; Rainey, 1989; Cheke et al., 1990; Chapman et al., 2004a; Reynolds et al., 2006; Maiga et al., 2008; Dingle, 2014), public health (Garms et al., 1979; Sellers, 1980; Ming et al., 1993; Ritchie & Rochester, 2001; Eagles et al., 2013; Huestis et al., 2019), and ecosystem vigor (Green, 2011; Landry & Parrott, 2016). We follow the definition of migration (sometimes considered as dispersal) as persistent movements that are unaffected by immediate cues for food, reproduction, or shelter, that have a high probability to relocate the animal in a distinct environment (Dingle & Drake, 2007; Dingle, 2014; Chapman et al., 2015). Recent radar studies have revealed the immense magnitude of insect migration, highlighting its role in ecosystem biogeochemistry via the transfer of micronutrients by trillions of insects moving annually in Europe (Hu et al., 2016; Wotton et al., 2019). Yet, these studies seldom provide species-level information (Drake & Reynolds, 2012), which is needed to discern the adaptive strategies, drivers, processes, and impacts of long-distance migration. Ideally, addressing these issues requires tracking individual insects over tens and hundreds of kilometers, a task that remains beyond reach for most entomological investigations due to the insects’ small size, speed, and flight altitudes that often exceed hundred meters above ground level (agl). Over the past decades, impressive advances in understanding of migration of a handful of large insects (>40 mg) provided key insights into migratory routes and the underlying physiology and ecology of migration with implications ranging from pest control to conservation (Chapman *et al*., 2008, 2012; Stefanescu *et al*., 2013; Hallworth *et al*., 2018). However, the identity of the vast majority of insect species engaging in migration remain unknown (Holland, 2006; Hu et al., 2016). With its growing human population and its nutritional demands, public health problems, and conservation challenges, such knowledge can be especially valuable for sub-Saharan Africa.

A longitudinal, systematic, comparative study on the high-altitude migration of multiple species of insects in the same region would be incredibly useful to identify variation in migration strategies and their ecological context, and help address some outstanding questions: Which species are the dominant windborne migrants? Do species migrate regularly or only in particular years? Within a year, does migration occur rarely, under specific weather conditions, or throughout the season? Do species exhibit uni-, bi-, or multi-directional flights and does the direction change with the season? How selective are the migrants for the wind direction they are harnessing? Using sticky nets mounted between 40 and 290m agl on tethered helium balloons, we undertook a three-year survey of flying insects in central Mali. Here, focusing on a dozen collected insect species—representing broad phylogenetic groups and ecological “guilds”—we seek to discern common migration patterns and infer underlying strategies. We characterized variation in migration over time and space to determine the regularity of flights in each species. Based on the phenology of the migrations and their relationship with meteorological conditions, including wind direction, which is a proxy of migration trajectory, we propose a simple classification of migration strategies of Sahelian taxa. We hope other investigators with an interest in the ecology and taxonomy of Afrotropical insects—especially pest species and their natural enemies—will utilize our aerial collections, and that these results will stimulate further collaborative research.

## Material and Methods

### Study area

Aerial sampling stations were placed in four Sahelian villages (Fig. 1): Thierola (13.6586, - 7.2147) and Siguima (14.1676, −7.2279; March 2013 to November 2015); Markabougou (13.9144, −6.3438; June 2013 to June 2015), and Dallowere (13.6158, −7.0369; July to November 2015). The study area has been described in detail previously (Lehmann et al., 2010; Huestis et al., 2012; Yaro et al., 2012; Dao et al., 2014; Lehmann et al., 2017). Briefly, it is a rural area characterized by scattered villages, with traditional mud-brick houses, surrounded by fields. A single growing season (June–October) allows farming of millet, sorghum, maize, and groundnuts, as well as small vegetable gardens. Over 90% of the rains fall during this season (~550 mm annually). Cattle, sheep, and goats graze in the dry savannah that consists of grasses, shrubs, and scattered trees. The dry and the wet seasons correspond to December–May and June–October, respectively, with >80% of the rainfall in July– September, forming puddles and seasonal ponds that usually dry by November. Rainfall is negligible from December until May (total precipitation 0–30 mm), when water is available only from deep wells.

**Figure 1.**
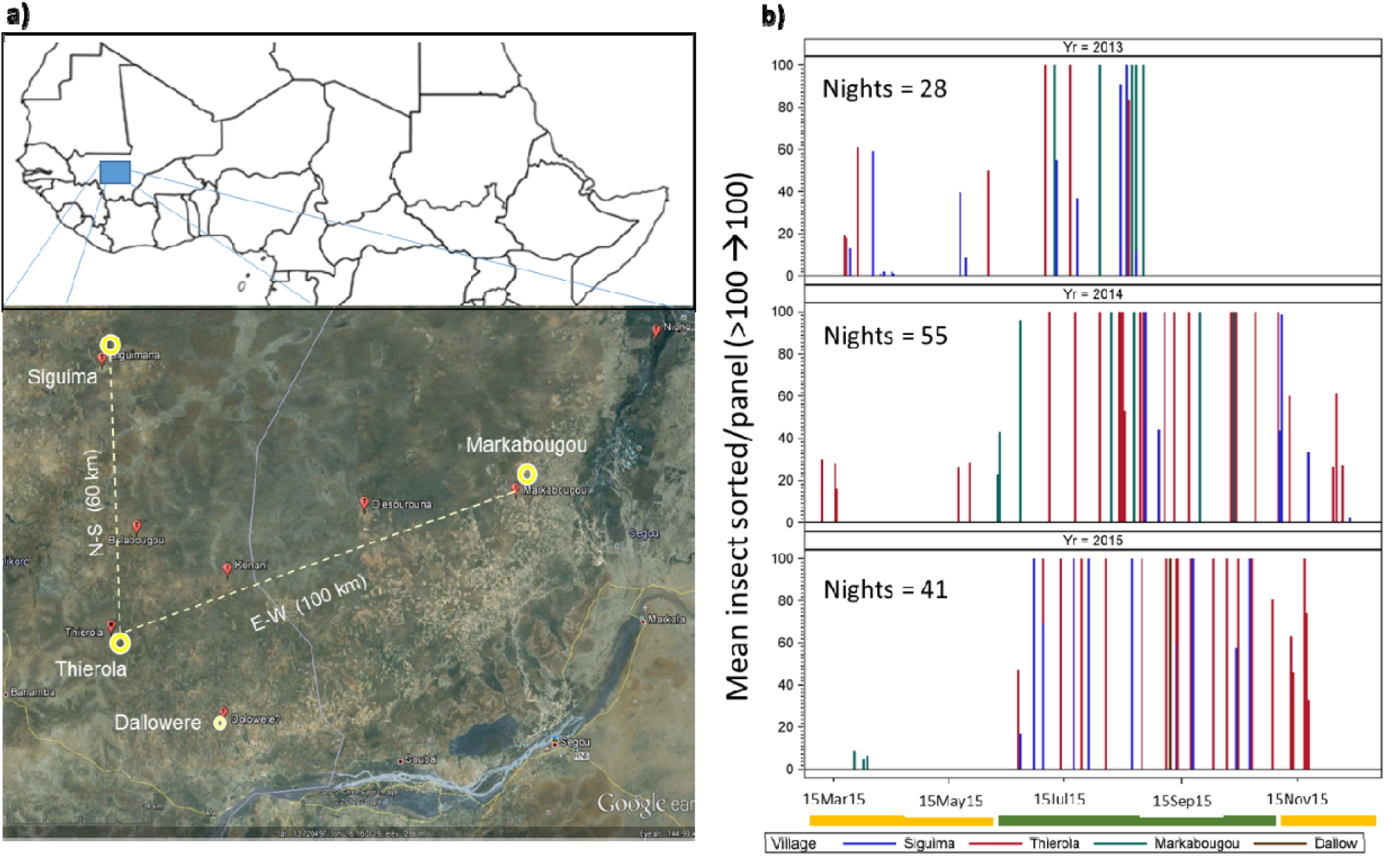
Sampling effort of high-altitude flying insects by year. Needles represent sampling nights (by village: color) and they extend up to 100 insects per panel (actual number of insects can exceed 2,000).

### Aerial sampling and specimen processing

The aerial sampling methods have been described in detail in Huestis et al. (2019), and are only briefly outlined here. Insect sampling was conducted using sticky nets (panels) attached to the tethering line of 3m diameter helium-filled balloons, with each balloon typically carrying three sticky nets. Initially, nets were suspended at 40m, 120m, and 160m agl, but from August 2013, the typical altitude was increased to 90m, 120m, and 190 m agl. When a larger balloon (3.3m dia.) was deployed at Thierola (August-September 2015), two additional nets were added at 240m and 290m agl. Balloons were launched approximately one hour before sunset (~17:00) and retrieved one hour after sunrise (~07:30), the following morning. To control for insects trapped near the ground as the nets were raised and lowered, comparable sticky nets (control panels) were raised up to 40m agl and immediately retrieved during each balloon launch and retrieval operation. Between September and November 2014, the control panels were raised to 120m agl. The control panels typically spent 5 minutes above 20m when raised to 40m, and up to 10 minutes when raised to 120m. Following panel retrieval, inspection for insects was conducted in a dedicated clean area. Individual insects were visually removed from the nets with forceps, counted, and stored in labeled vials containing 80% ethanol.

### Taxon selection and identification

Using a dissecting microscope, insects were sorted by morphotype—an informal taxon assigned to specimens with similar morphology that are putative members of a single species— counted and recorded in a database. The remaining insects were sorted to order, counted, and again recorded. Selected morphotypes were chosen based on their easily identifiable features and their repeated appearance in a preliminary examination of the collection. Later, a subset (N>10) were identified by expert taxonomists who narrowed the identification down to species or genus and confirmed that the morphotype likely represents a single species (Tables S1 and S2). The thirteen taxa used in the present study are described in Table S2 and Fig. S1.

### Statistical Analysis

From each site we aimed to sample 2–3 dates of aerial collection per sampling month. When aerial sampling was done in only one or two villages up to 4 dates of collection were selected per month. The dates were spread more or less evenly through the sampling days of each month. From each sampling night, two nets were selected and 1–4 vials of insect specimens, aiming to represent >30% of the total insects collected, were sorted and counted as described above. To compute the ‘panel density’ of the selected taxa, the product of the total number of insects collected on that panel and the fraction of specimens from each taxon in the subsample sorted was rounded to the nearest integer.

The Modern-Era Retrospective analysis for Research and Applications, Version 2 (<u>MERRA-2</u>) atmospheric reanalysis fields (Gelaro et al., 2017)—selected to represent observed nightly conditions (18:00 through 06:00)—were used to calculate nightly mean temperature, relative humidity (RH), wind speed and direction. Corresponding values were computed for 2, 50, 70, 200, 330m agl for the nearest grid center (available in ~65km^2^ resolution) of each village: Siguima, Markabougou and Thierola (Dallowere, located 25km south of Thierola was included in the same grid of Thierola). Temperature, RH, wind speed and direction were available as hourly up to 10m, and as 3-hourly at altitude >10m. Conditions at panel height, e.g., mean nightly wind speed, were estimated based on the nearest available altitude layer. Mean nightly wind direction was computed based on the average values of north-south and east-west vectors of the hourly wind direction angle (unweighted by wind speed). The altitudes of the sampling panels were generally well above the insects’ ‘flight boundary layer’—the lowest air layer where an insect’s self-propelled flight speed is greater than wind speed—so flight direction is governed primarily by wind direction (Taylor, 1974; Chapman et al., 2011; Drake & Reynolds, 2012). Thus, flight direction was estimated from the average nightly wind direction during the night of capture at the location and panel height. The seasonal flight direction was estimated as the weighted average nightly wind direction at the flight altitude at which a taxon was captured, with the taxon aerial density at the panel used as a weight to compute the weighted circular mean angle for that taxon (see above). The dispersion of individual angles around the mean was measured by the mean circular resultant length ‘r’ (range: 0 to 1), indicating tighter clustering around the mean by higher values. Rayleigh’s test of uniformity was used to test whether there was no mean direction, as when the angles form a uniform distribution over 360 degrees. Data analysis was carried out both on panel (net) density, and aerial density, to ensure that all key aspects of the data are well represented.

Aerial insect density was estimated based on the taxon’s panel density (above) and total air volume that passed through that panel that night, i.e.:

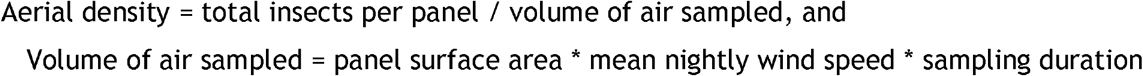

Insect sampling duration was calculated from balloon launch time until its retrieval time (typically 17:30 to 7:30; ~14h). Based on panel altitude, wind speed was selected from the nearest layer (above) to calculate the nightly average for each panel. This calculation assumed that air passed through the net with minimal attenuation and that the panel remained perpendicular to the wind direction throughout. These are reasonable assumptions because observations showed no substantial tilting or flipping of the nets, the thin layer of glue did not block the holes of the net, and insects were always found only on one side of the net. Under these assumptions (panel surface = 3m^2^, for 14h) and nightly average wind speed from the MERRA-2 database, the volume of air that would pass through the sticky nets when average nightly wind speed was 1 vs. 7m/s is 151,200 and 1,058,400m^3^ of air, respectively.

Data analysis was initially carried out on the panel density for a given taxon to provide a direct description of the main trends. Thereafter, a comprehensive statistical analysis was carried out on corresponding aerial density. The variation in the aerial density measured by each panel due to the effects of season, panel height, and other factors was evaluated using mixed linear models with either *Poisson* or negative binomial error distributions implemented by proc GLIMMIX with a log link function (SAS Inc., 2011). These models accommodate the non-negative integer-counts and the combination of random and fixed effects. The ratio of the Pearson χ^2^ to the degrees of freedom was used to assess overall “goodness of fit” of the model, with values greater than two indicating poor fit. The significance of the scale parameter estimating k of the negative binomial distribution was also used to choose between *Poisson* and negative binomial models. Lower Bayesian Information Criterion (BIC) values and the significance of the underlying factors were also used to select the best fitting model of each taxon.

## Results

In total, 461,100 insects were collected on 1,894 panels between 2013 and 2015. Sorting of 4,824 specimens representing 77 panels from altitudes between 40m and 290m agl revealed a diverse assembly representing thirteen orders (Fig. 2a and Table 1). Members of the Coleoptera dominated these collections at 53%, followed by Hemiptera (27%) – especially Auchenorrhyncha (18.5%), Diptera (11%), and Hymenoptera and Lepidoptera at 4% each, together accounting for >99% of the insects collected (Table 1). Additional specimens were identified totaling 100 insect families (Tables 1 and S1). Family, genus, and species level identification of the insects was carried out by a dozen participating taxonomists, exclusively on their group of expertise, thus we are confident that many other species and families are yet to be recognized in these collections. To explore patterns of high-altitude flights of Sahelian insects, thirteen species representing diverse phylogenetic and ecological groups were identified and counted from samples consisting of a total of 25,188 high-altitude flying insects. These represent additional subsamples totaling 58,706 insects captured on 222 nets over 125 aerial collections, carried out over 96 sampling nights between 2013 and 2015, in one or more of the Sahelian villages (Fig. 1).

**Table 1.** Overall diversity of insects collected in aerial samples (40-290 m agl) as reflected by insect order composition (see also Figure 2). The insect families sent for identification by taxonomists represent a small fraction of the predicted total diversity (see text for details). Orders represented by 1-2 specimens (Blattodea, Thysanoptera, Megaloptera, Psocoptera, and Phasmatodea) are not shown.

| Order | Percent | Families identified | Families |
| --- | --- | --- | --- |
| Coleoptera | 53.3 | Aderidae, Anthicidae, Attelabidae, Bostrichidae, Brentidae, Carabidae, Chrysomelidae, Coccinellidae, Curculionidae, Dytiscidae, Elateridae, Erorhinidae, Hydrophilidae, Mordellidae, Nitidulidae, Phalacridae, Scarabaeidae, Staphylinidae | 18 |
| Hemiptera<br>(Heteroptera) | 8.2 | Berytidae, Corixidae, Cydnidae, Geocoridae, Gerridae, Hydrometridae, Lygaeidae, Miridae, Nabidae, Notonectidae, Oxycarenidae, Pentatomidae, Pyrrhocoridae, Reduviidae, Rhopalidae, Rhyparochromidae, Stenocephalidae, Tingidae, Veliidae | 19 |
| Hemiptera<br>(Homoptera) | 19 | Aphididae, Cicadellidae, Delphacidae, Flatidae, Ricaniidae | 5 |
| Diptera | 11.2 | Anthomyiidae, Calliphoridae, Cecidomyiidae<sup>b</sup>, Ceratopogonidae<sup>b</sup>, Chironomidae<sup>b</sup>, Chloropidae, Culicidae<sup>b</sup>, Curtonotidae, Diopsidae, Dolichopodidae, Drosophilidae, Ephydriidae, Lauxaniidae, Limoniidae<sup>b</sup>, Lonchaeidae, Milichiidae, Muscidae, Mycetophilidae<sup>b</sup>, Phoridae, Pipunculidae, Platystomatidae, Rhiniidae, Sepsidae, Simuliidae, Tachinidae, Tephritidae, Tipulidae, Ulidiidae | 28 |
| Hymenoptera | 4 | Apidae, Bethylidae, Brachonidae, Chalcididae, Chrysididae, Crabronidae, Diapriidae, Dryinidae, Eulophidae, Eupelmidae, Eurytomidae, Figitidae, Formicidae, Ichneumonidae, Megachilidae, Pompilidae, Rhopalosomatidae, Scelionidae, Sphecidae | 19 |
| Lepidoptera | 3.9 | Gelechiidae<sup>b</sup>, Nolidae<sup>b</sup> | 2 |
| Orthoptera | 0.3 | Acrididae, Gryllidae, Pyrgomorphidae, Tetrigidae, Tettigonidae, Trigonidiidae | 6 |
| Neuroptera | 0.2 | Chrysopidae, Mantispidae, Myrmeleontidae | 3 |
| Total |  |  | 100 |

<sup>a</sup> In total, 4,824 insects from 77 sticky nets (panels) were used to estimate the order composition.
<sup>b</sup> Identified through DNA barcoding correlations by Dr. Yvonne Lint

**Figure 2.**
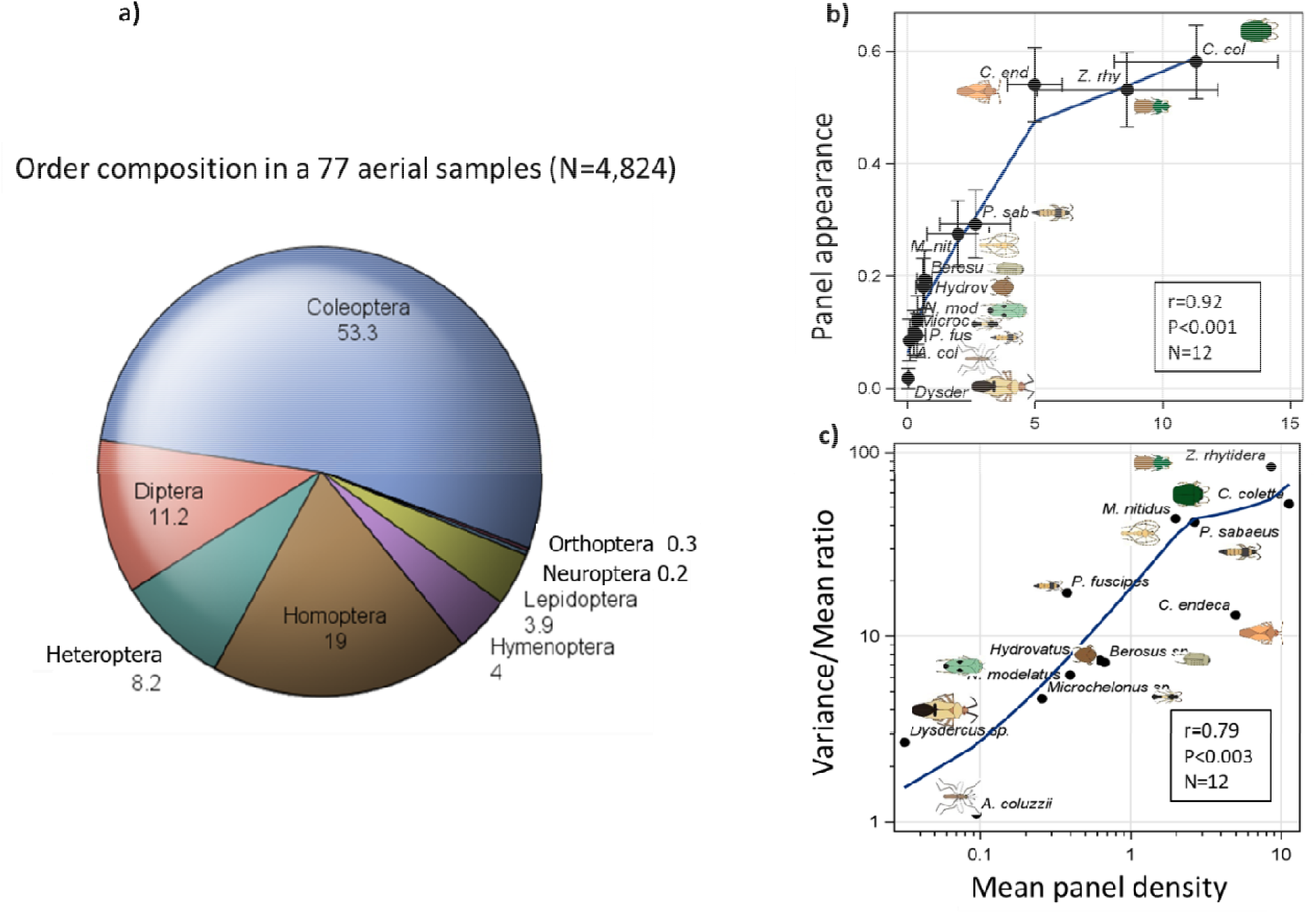
a) Overall diversity (by insect orders) of aerial collection estimated based on samples from 70 sticky nets. Orders represented by less than 3 specimens (Blattodea, Thysanoptera, Megaloptera, Psocoptera, and Phasmatodea) were not shown. b) The relationship between the variance to mean ratio and its mean panel density. c) Relationship between overall species density/panel (+95% CI) and the fraction of nets it was captured on (+95% CI) as a measure of the regularity of HAF activity. Insects show the Pearson correlation coefficient (r), its P value (P) and sample size (N). Schematic insect silhouettes are not to scale.

In the test to ascertain that insects were collected at high altitude and not inadvertently trapped near the ground as the nets were raised and lowered, it was determined that only 564 insects were captured on 507 control panels compared with 58,706 captured on the 222 experimental (standard) panels. The control panels spent 3–5 minutes above 30m and thus could be expected to contain ~0.5% of a normal panel that remained at high altitude for 14 hours, assuming aerial density of the insects remained constant over that time. To assess if insects were intercepted below 30m agl, we tested if the mean panel density of each taxon on the control panel was (i) significantly lower than corresponding mean on standard panel, and (ii) that it was not significantly higher than the equivalent of a 4-minute aerial nightly sampling with a standard panel (Table 2). Except for the *Hypotrigona* sp. (Hymenoptera), both expectations were met for all taxa, with a control to standard mean density ratio of 0– 0.003. Such a low ratio suggests that most insects engage in high-altitude flight during part of, rather than the whole night and that this part does not include the launch or retrieval times. For the *Hypotrigona* sp., however, the ratio of mean control to standard panel was 0.167, suggesting that ~17% of its aerial density could have been collected near the ground (Table 2). For this reason, this taxon was removed from further analysis.

**Table 2.** Overall abundance and occurrence of selected taxa in aerial samples collected on standard panels (220 panels between 40 and 290 m agl, in 125 sampling nights) and control panels (508 nets between 40 and 120 m agl)

| ----- Standard panels (222 panels in 125 sampling nights) ----- |  |  |  |  |  |  |  |  |  | ----- Control panels (507 panels) ----- |  |  |  |  |  |  |  |
| --- | --- | --- | --- | --- | --- | --- | --- | --- | --- | --- | --- | --- | --- | --- | --- | --- | --- |
| Taxon | Mean panel density | Lower 95% CI<sup>a</sup> | Median panel density | Panel Freq<sup>b</sup> | Night Freq<sup>c</sup> | Total | Max panel density | Var / mean | Mean contr / standr<sup>d</sup> | Mean panel density | Upper 95% CI<sup>e</sup> | Four min Standr<sup>f</sup> | Med panel density | Panel Freq<sup>b</sup> | Total | Max panel density | Var / mean |
| *Dysdercus* sp. | 0.03 | -0.007 | 0 | 0.02 | 0.04 | 7 | 4 | 2.7 | 0.0000 | 0 | nd | 0.0001 | 0 | 0.0000 | 0 | 0 | nd |
| *Cy. endeca* | 5.00 | 3.9325 | 1 | 0.54 | 0.67 | 1110 | 44 | 13.0 | 0.0016 | 0.0079 | 0.0156 | 0.0230 | 0 | 0.0079 | 4 | 1 | 0.993 |
| *M. nitidus* | 1.99 | 0.7636 | 0 | 0.27 | 0.43 | 442 | 117 | 43.2 | 0.0010 | 0.002 | 0.0058 | 0.0092 | 0 | 0.0020 | 1 | 1 | 1 |
| *N. modulatus* | 0.40 | 0.1898 | 0 | 0.12 | 0.21 | 88 | 17 | 6.2 | 0.0000 | 0 | nd | 0.0018 | 0 | 0.0000 | 0 | 0 | nd |
| *A. coluzzii* | 0.09 | 0.0519 | 0 | 0.09 | 0.17 | 21 | 2 | 1.1 | 0.0000 | 0 | nd | 0.0004 | 0 | 0.0000 | 0 | 0 | nd |
| *P. sabaesus* | 2.66 | 1.2729 | 0 | 0.29 | 0.42 | 591 | 134 | 41.4 | 0.0000 | 0 | nd | 0.0122 | 0 | 0.0000 | 0 | 0 | nd |
| *P. fuscipes* | 0.38 | 0.0414 | 0 | 0.09 | 0.21 | 84 | 32 | 17.2 | 0.0000 | 0 | nd | 0.0017 | 0 | 0.0000 | 0 | 0 | nd |
| *Z. rhytidera* | 8.61 | 5.0629 | 1 | 0.53 | 0.61 | 1912 | 246 | 83.6 | 0.0000 | 0 | nd | 0.0396 | 0 | 0.0000 | 0 | 0 | nd |
| *Ch. coletta* | 11.31 | 8.1055 | 2 | 0.58 | 0.71 | 2511 | 174 | 51.9 | 0.0009 | 0.0099 | 0.0185 | 0.0520 | 0 | 0.0099 | 5 | 1 | 0.991 |
| *Hydrovatus* sp. | 0.63 | 0.3414 | 0 | 0.18 | 0.34 | 139 | 21 | 7.4 | 0.0000 | 0 | nd | 0.0029 | 0 | 0.0000 | 0 | 0 | nd |
| *Berosus* sp. | 0.67 | 0.3805 | 0 | 0.19 | 0.32 | 149 | 20 | 7.2 | 0.0030 | 0.002 | 0.0058 | 0.0031 | 0 | 0.0020 | 1 | 1 | 1 |
| *Microchelonus* sp. | 0.26 | 0.1132 | 0 | 0.10 | 0.19 | 57 | 11 | 4.6 | 0.0000 | 0 | nd | 0.0012 | 0 | 0.0000 | 0 | 0 | nd |
| *Hypotrigona* sp. | 0.19 | 0.0519 | 0 | 0.05 | 0.08 | 42 | 8 | 5.7 | 0.1670 | 0.0316 | 0.0754 | 0.0009 | 0 | 0.0316 | 16 | 11 | 7.982 |
| Overall | 2.48 |  |  | 0.24 | 0.34 | 7153 |  | 21.9 | 0.0005 | 0.0018 |  |  |  | 0.0018 | 27 |  | 0.996 |

<sup>a</sup> The lower 95% confidence interval (CI) of the mean panel density of standard panel used to compare with the upper 95% CI of the control panel for each taxon (see text).
<sup>b</sup> Panel Freq - Frequency of panels with at least one specimen per taxon.
<sup>c</sup> Night Freq - Frequency of nights with at least one specimen per taxon per night regardless of village (ie., includes nights when launches occurred in more than one village, n = 96).
<sup>d</sup> Overall mean ratio of control/standard panel and mean control panel density were computed excluding *Hypotrigona* sp. (see text).
<sup>e</sup> Expected density assuming insects were intercepted while the control panels reached over 20m and remain there for ~four minutes (see text).

### Overall abundance, sampling distribution, and correlation between taxa

Overall abundance of the selected taxa, measured as mean panel density ranged over two orders of magnitude, with lowest mean panel density near 0.05 for the *Dysdercus* sp. (Hemiptera) and *Anopheles coluzzii* (Diptera) compared to 11 for *Chaetocnema coletta* (Coleoptera, Table 2, Fig. 2b). The distribution of captured insects on (standard) nets was strongly “L shaped,” typical of clumped distributions (Fig. S2), with a median panel density of zero for all species except *Ch. coletta*, <u>*Cysteochila endeca*</u> (Hemiptera) and *Zolotarevskyella rhytidera* (Coleoptera, Table 2), and a maximum of 246 specimens per panel, raising the possibility that flight activity was concentrated on one or a few nights. The nightly occurrence frequency varied between 4 and 71% (Table 2), indicating that all taxa engaged in high-altitude flight activity over multiple nights. Moreover, panel occurrence frequency was positively correlated with taxon panel density (r_P_=0.92, P<0.001, N=12, Fig 2b), indicating that taxa that appeared on fewer nights were the least abundant and that their low overall abundance may account for their low appearance. Nonetheless, the high values of the variance to mean ratios (2.7–83.6, Table 2 and Fig. 2c) of all taxa, except *A. coluzzii*, suggest that the distribution of insects was temporally clustered.

All taxa exhibited marked seasonality in high-altitude flight activity (Fig. 3a), peaking between July and October, following a considerably lower activity in May–June. Overall, flight declined substantially in November–December, and virtually none was recorded in March-April; presumably this was also the case earlier in the dry season (January and February), when no aerial samples were taken (Fig. 1). Visual examination of the seasonality of individual taxa (Fig. 3a) suggests variability among species in flight activity. For example, *Microchelonus* sp. (Hymenoptera) appeared as early as May and peaked in June, whereas, the leafhopper, *Nephotettix modulatus* (Hemiptera) first appeared in July and peaked in October (Fig. 3a). A unimodal activity best describes *A. coluzzii*, *Dysdercus* sp., *Microchelonus* sp., and *N. modulatus*, while bimodal activity describes the other taxa, e.g., *Paederus fuscipes* (Coleoptera), and *Berosus* sp. (Coleoptera, Fig. 3a). Bimodal distribution was also suggested by the total-insect density/panel (Fig. S3) Likewise, correlations between taxa in nightly flight were low (Spearman, r_S_ mean=0.17, −0.15<r_s_<0.62, Fig. 3b). The highest r_S_ -values involved high density taxa, e.g., *Paederus sabaeus* and *Metacanthus nitidus* (r_s_=0.62). The mean pairwise correlation dropped to 0.09 after excluding December through April, indicating that the dry season, when migration is negligible, contributed substantially to the overall correlation between taxa. Low positive correlation indicates species-specific dynamics of migration concentrated around the wet season, rather than rare mass migration events.

**Figure 3.**
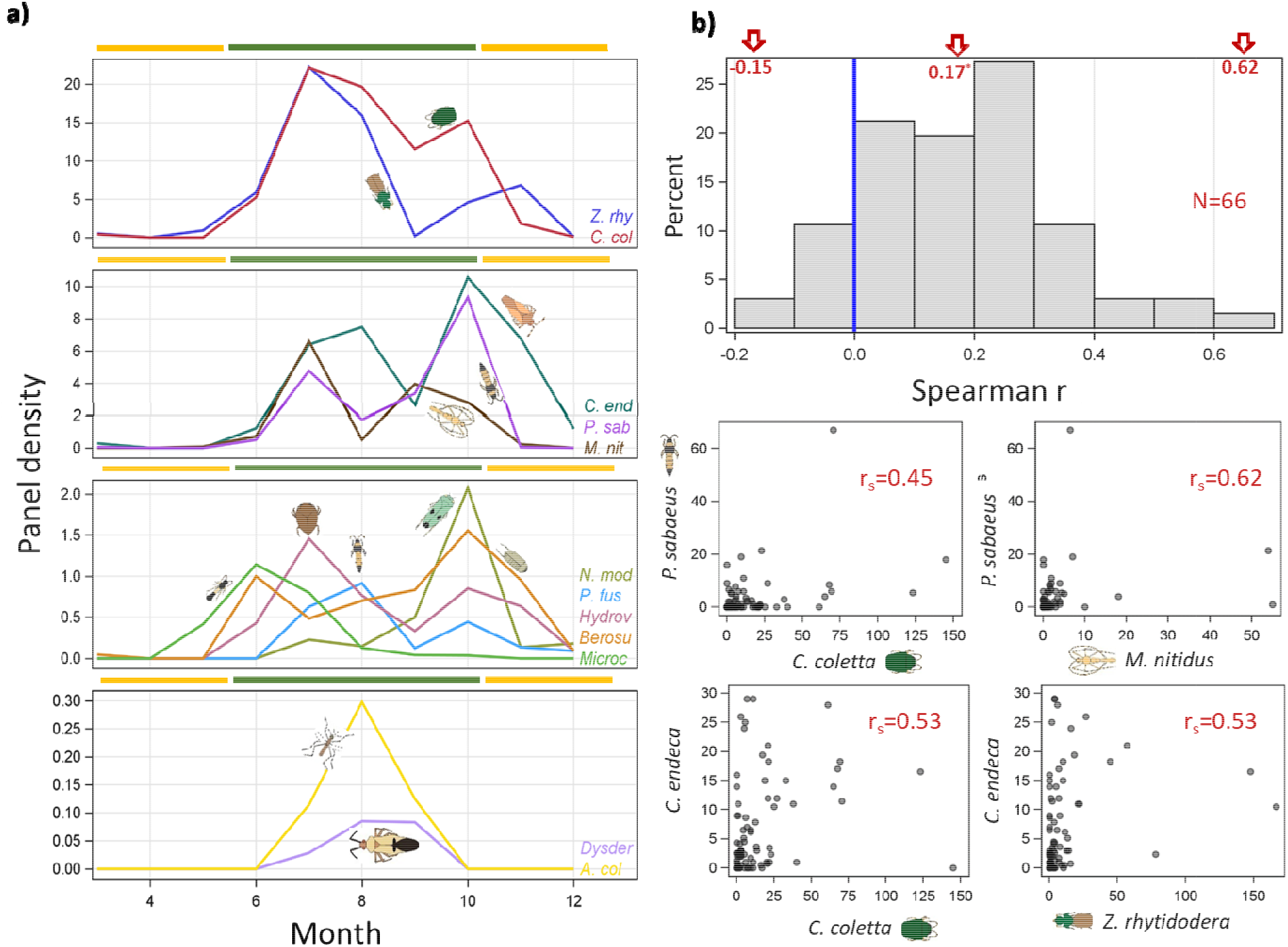
Temporal variation in flight activity across taxa. (a) Seasonal variation of migrant insects measured by panel density based on three-year data. Dry and rainy season are shown by yellow and green colors (ruler). (b) The distribution of the Spearman correlation coefficient (r_s_) between 66 pairs of migrant insects and relationship in nightly mean densities of the taxa pairs with highest Spearman correlation coefficients (b, N=95 nights). Schematic insect silhouettes are not to scale (species names are truncated to conserve space).

### Variation between collection sites, years and altitude

The fraction of positive appearances of each taxon was compared between localities (up to 100 km apart) and years to assess if they were location- or year-specific (Fig. 4). All taxa were found in all locations (Dallowere, situated 25 km from Thierola was excluded from the comparison because it was represented by only two sampling nights). The similarity between the localities in the appearance of each taxon is striking since they were partly sampled in different dates and even different months (Fig. 1). Similarly, all taxa were present in every sampling year except for *Dysdercus* sp. and *N. modulatus*, which were not sampled in 2013. The comparative rarity of *Dysdercus* sp. may account for its absence from the sparse data in 2013, which consisted of 28 sampling nights vs. 41 and 56 in the other villages. Likewise, *N. modulatus* appeared late in the season (peaks in October) during 2014 and 2015, thus, it was unlikely to be sampled in 2013, in which collection ended by mid-August.

**Figure 4:**
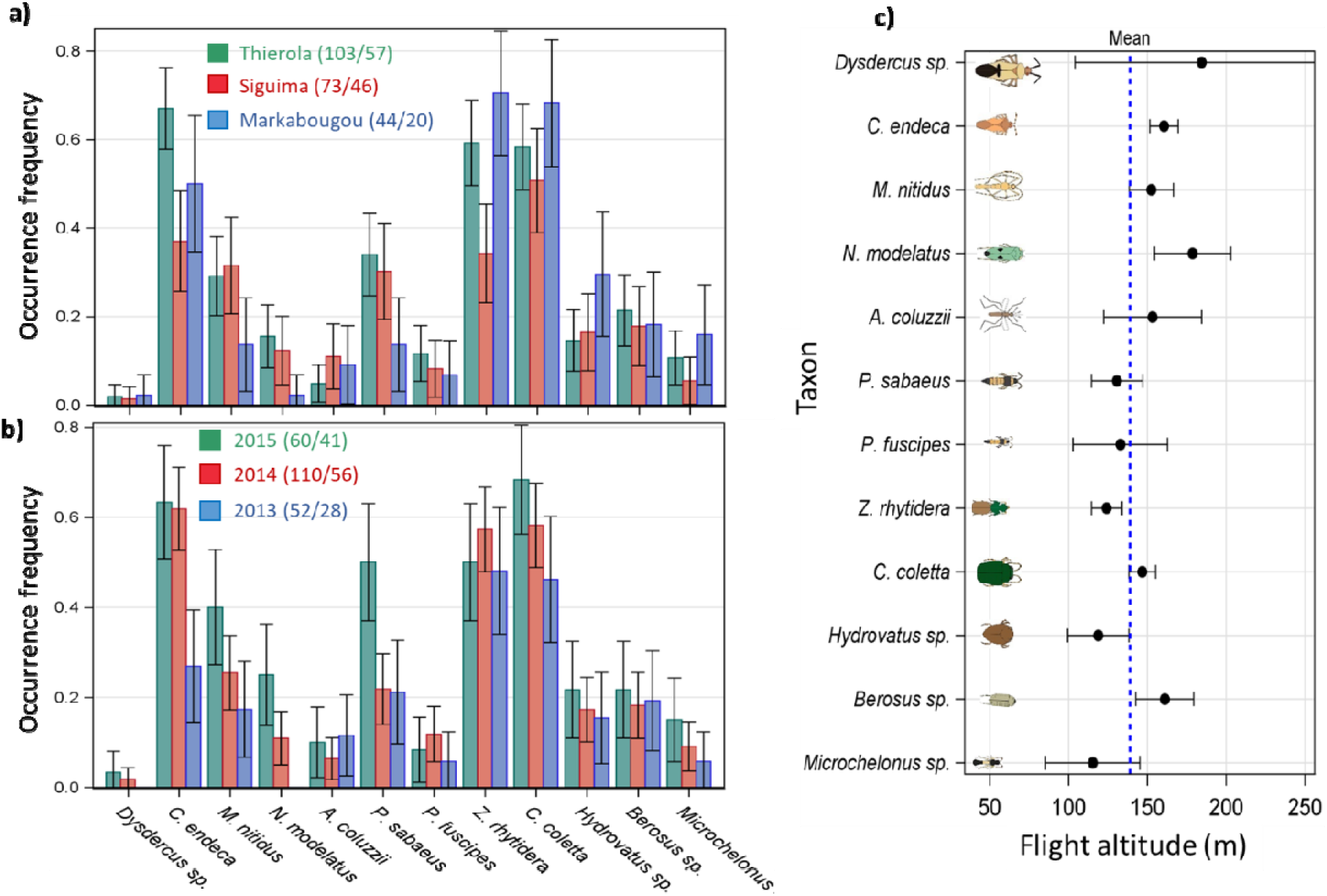
Spatial and annual variation in high altitude migration. Mean frequency of occurrence (+ 95% CI) of each taxon per panel by (a) locality (excluding Dallowere which was sampled in only 2 nights) and (b) year. The sampling effort in each year with respect to nets and nights is given in the legend. Between-species variation in flight altitude measured as mean panel altitude (+95% CI) weighted by panel density (c). Dotted blue line shows mean panel altitude. Note: the highest panel was typically 190 m agl, but between August and September 2015 we used a larger helium balloon and the highest panel was set at 290 m agl. (see Methods). Schematic insect silhouettes are not to scale.

The typical flight altitude of each taxon, measured as an average panel height weighted by the taxon’s panel density values, varied among taxa from 130m (*Microchelonus* sp.) to 175m (*N. modulatus*, Fig. 4c). Despite limited use of the largest balloon (3.3m in diameter), which allowed sampling up to 290m agl in Thierola between August–September 2015, all taxa were collected in the top nets (240–290m agl), suggesting that our sampling has not reached their highest flight altitude, in accordance with radar data suggesting that migrants reach (and often exceed) 450m agl (Reynolds & Riley, 1988; Reynolds et al., 2005; Wood et al., 2009; Drake & Reynolds, 2012). Likewise, all taxa were captured in the lowest panel (40m agl), except *Dysdercus* sp. and *N. modulatus*, which were not captured below 120m agl. The low abundance of *Dysdercus* sp. may be accounted for by incomplete altitude sampling, but presumably *N. modulatus* tends to fly at higher altitudes. Significant differences between species suggest that low-flying insects including *P. sabaeus*, *Z. rhytidera*, and *Hydrovatus* sp. may engage in shorter flights than high-flying taxa, e.g., *N. modulatus*, *Cy. endeca*, *Berosus* sp., and *M. nitidus*.

### Aerial density and the effects of weather on high-altitude migration

Estimated aerial density of each taxon, expressed as density/10^6^ m^3^ of air (Methods), was positively correlated with panel density (r = 0.93, P<0.001, Fig. S3a), suggesting that the results of analyses based on panel- and aerial-density would be similar. This high correlation stemmed in part from the modest variability in average nightly wind speed at flight altitude (mean = 5.1m/s, and the 10^th^ and 90^th^ percentiles are 2.4 and 8.0m/s, respectively, Fig S4).

The aerial density values were more clustered than panel density values based on the variance to mean ratio (Tables 2 and 3). Even after accounting for the variation due to season, year, village, and altitude, models with a negative binomial error distribution were superior to those with *Poisson* distributions in all taxa (Table 3). Similar to the results based on panel density (Fig. 4), variation due to year and locality (village) of sampling were not significant in all taxa, whilst seasonality was significant in seven taxa (P<0.05, Table 3 and Fig. 3a), as was the case for altitude. However, the effect of altitude was modest (Fig. 4c). In subsequent models, the effects of village and year were therefore removed.

**Table 3.**
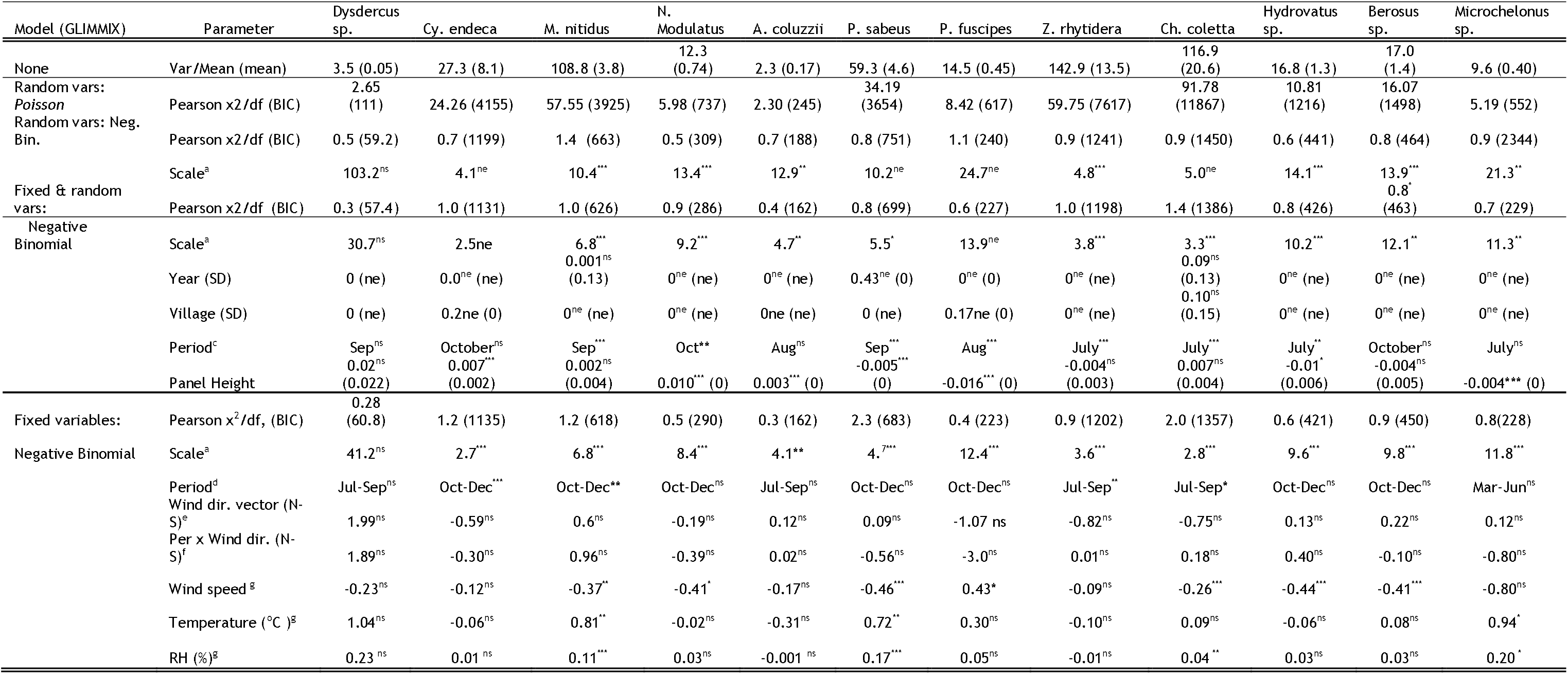
Variation in taxon’s aerial density among years, locality (villages), altitude, and meteorological conditions (GLIMMIX models of random (year and village) and fixed (season, panel height, wind speed and direction, temperature and RH at flight height, 222 nets between 40 and 290 m agl, in 125 sampling nights).

The typical weather conditions near the ground (2 m agl) and at flight-height, expressed as the mean values weighted by the corresponding aerial density of each taxon varied modestly between taxa as reflected by the many overlapping 95% CI of different taxa (Fig. 5). The wet season mean nightly temperatures of the ground and air intersected the 95% CI of 8 and 9 of the taxa, respectively, suggesting that, during the wet season, temperature is unlikely to limit migration (Fig. 5a). A similar pattern held for RH and wind speed, although most insect flight activity took place at lower RH and lower wind speed than their wet season averages (Fig. 5), possibly because very rainy and stormy conditions inhibit migration (or because aerial sampling did not include such nights). Considering variation among taxa, the mosquito *A. coluzzii* exhibited peak high-altitude flight at lower temperatures (over ground and at flight altitude) and higher RH, whilst *Berosus* sp. and *Microchelonus* sp. exhibited peak high-altitude flight at higher (ground and flight) temperatures and lower RH (Fig. 5). This pattern may confound the seasonal activity because the rains are most frequent in August, when *A. coluzzii* exhibited its peak activity, whereas *Berosus* sp. and *Microchelonus* sp. peaked at October and June (Fig. 3) when rains are rare, and temperatures are higher while RH is lower. Statistical models that evaluated the effects of these weather parameters on high-altitude flight in the presence of period (March–June, July–September, and November–December) revealed that flights of certain taxa were more common under lower wind speed (seven taxa) and higher RH (four taxa, Table 3, see below about wind speed).

**Figure 5:**
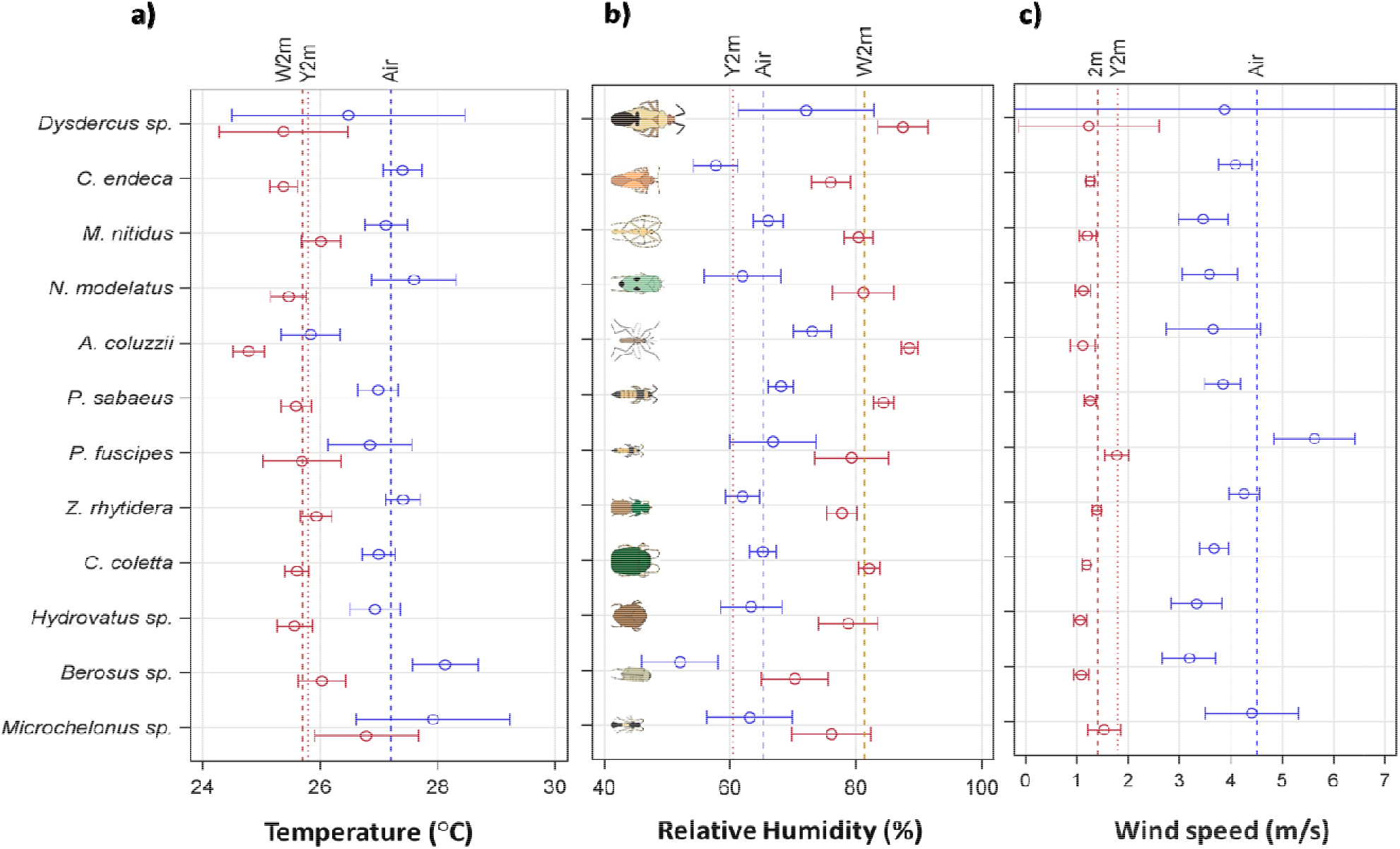
Weather conditions during high-altitude flight on the ground (red) and at flight height (blue) of each taxon. Weighted means (weighted by aerial density) and 95% CI on the ground and at flight height of (a) nightly temperature, (b) RH, and (c) wind speed are shown with corresponding wet season means (dashed lines labeled ‘W2m’ and ‘Air’) and year-round means at 2 m agl (dotted red lines labeled ‘Y2m’). Schematic insect silhouettes are not to scale.

### Seasonal wind and flight directions

As expected (Nicholson, 2013), wind direction in the Sahel showed marked seasonality, with northerly winds (blowing towards the south) dominating from December to April and reversing course from May to October (Fig. 6a). During these periods >70% of the winds could transport insects in the seasonally-beneficial “preferred” direction, i.e., northward during the wet season, and southward during the dry season. However, in November, winds are variable and blow towards the south and the north in similar frequencies (Fig. 6a).

**Figure 6:**
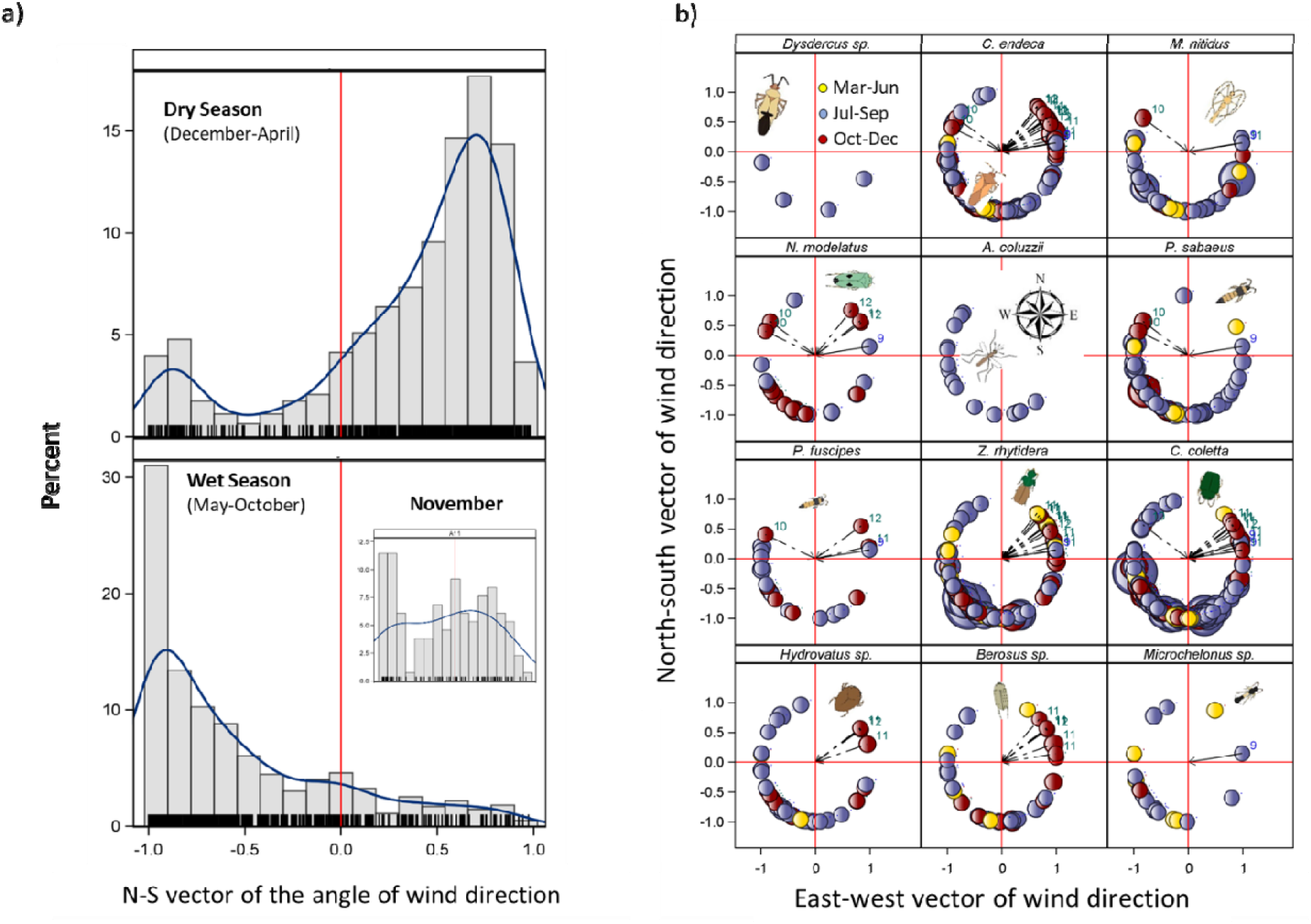
Seasonality of the south-north component of nightly wind direction in the Sahel and nightly wind direction during high-altitude flights of each taxon. a) To explore the possibility of north-south migration into the Sahel from more equatorial regions, the north-south component of nightly wind direction (2012-2015 MERRA2 data; all nights) shows the frequencies of winds during the dry (top) and wet (bottom) season in Thierola (the other villages exhibited similar distributions). Kernel distributions are shown in blue. Wind direction from the N and S are indicated by positive and negative S-N vector values, respectively. INSET: November is a transition month with variable wind direction. Red reference line at the origin indicates easterly or westerly winds. Fringe marks indicate actual values N-S component of wind direction. b) Wind direction during high-altitude flights of selected taxa. Circles denote source of mean nightly winds in relation to the capture location (origin) with north and east denoted by top and right red lines, respectively. Circle size reflects nightly aerial density and their color denotes the period (top left). Dotted arrows highlight southbound winds during the end of the wet season, that could be used for the “return” migration from the Sahel towards tropical areas closer to the Equator (numbers denote the months of such events). Schematic insect silhouettes are not to scale.

All taxa exhibited frequent northward migrations on southerly winds, which ranged widely from WSW to the ESE during the wet season (Fig. 6b). Such movements support the recolonization of the Sahel by “non-resident taxa” from broadly southern populations. During the wet season (July–September), Sahelian rainfall is associated with large mesoscale convective systems with squall lines which have changeable wind directions. So, insects could have been carried by winds to nearly all directions (Fig. 6) and dispersed widely across the Sahel, albeit with different intensities. The concentration of the circles, denoting the source of mean winds, and their size, which correspond to aerial density (Fig. 6b), likely signifies the relative position of the source populations. For example, *Z. rhytidera* exhibited a strong influx from the southwest, and *Ch. coletta* from western and southern sources (Fig. 6b). An approximately uniform distribution over large sectors, on the other hand, e.g., *Hydrovatus* sp. (Dytiscidae) and *Berosus* sp., may indicate dispersal with only weak concentration of southerly origin, probably reflecting migration between aquatic habitats in multiple directions.

To assess if a movement back into the savannas south of the Sahel took place at the end of the rainy season, we examined whether insects exhibited movement with southbound winds during September to December (Fig. 6b). Although they were less common, at least one or a few nights’ migration on southward wind were recorded in all taxa except *Dysdercus* sp., *A. coluzzii*, and *Microchelonus* sp. Notably, all three were the least abundant taxa (Fig 2c). Such southward movements were especially frequent in *Z. rhytidera*, *Cy. endeca*, *Ch. coletta*, and *Berosus* sp. (Fig. 6b). Tests of wind “selectivity” performed using contingency tables contrasting the proportion of nights with northbound and southbound flights during October through December (as well as October–November), exhibited no support for selective southward flight across taxa (P>0.05 at the individual test level, not shown). Similarly, no significant interaction of wind direction by period was detected in the aerial density analysis (Table 3).

## Discussion

This study presents the first cross-season survey of high-altitude migrant insects in Africa. Based on 125 high-altitude sampling nights, totaling 1,800 sampling hours, and yielding 222 samples from individual nets (and 507 additional control nets), we assessed the diversity of migrants and focusing on a dozen taxa, evaluated their flight regularity, seasonality, direction, and relationship with key meteorological conditions. Comparison between species may allow discerning common patterns that might fit other species in this region or beyond, leading to better understanding of these adaptive strategies.

The large number of taxa already identified from a small fraction of the aerial collection (Table 1 and Supplementary Table 1) suggests that migration at altitude is a common and widespread life history strategy in the Sahel, as expected from the impermanence of many habitats (Southwood, 1962; Drake & Gatehouse, 1995). Dominated by Coleoptera (53%), Hemiptera (27%, especially *Auchenorrhyncha*), and Diptera (11%, Table 1), the composition of this collection was distinct from those reported in Europe—dominated by Hemiptera (especially aphids) and Hymenoptera (Chapman et al., 2004b)—and in North America (Glick, 1939) —dominated by Diptera and Coleoptera, possibly reflecting taxa more tolerant of xeric environments. Insects flying >200m agl and day-flyers were underrepresented in our collection, as were larger insects (e.g., grasshoppers, moths) that could detach themselves from the thin layer of glue. Also, tiny insects (e.g., aphids and midges) might have been overlooked when insects were manually extracted from the nets. Nonetheless, a subsample of less than 10% of our aerial collections revealed members of 13 orders (Table 1) and, although only a handful were identified to the family level, >100 families were represented (Tables 1 and S1). Among the insect genera identified, several included known or suspected windborne migrants, such as *Dysdercus* sp. (Duviard, 1977), *A. coluzzii* (Garrett-Jones, 1962; Dao et al., 2014; Huestis et al., 2019), and *M. nitidus* (Morkel & Jacobs, 2014), but for the majority of the species, such knowledge is new, as is their reported presence in Mali. Clearly, this aerial collection awaits additional study and the authors would be pleased to hear from readers who might be interested in undertaking further identifications and study of particular taxa from this collection.

High-altitude migration occurred regularly over the three years of this study, peaking during the wet season (Figs. 3 and 4), suggesting that windborne migration is integral behavior in these taxa. The hypothesis that migration occurred exclusively on particular nights, suggested by values of the ratio of the variance to the mean (4.6–83.0) and the zero median panel density for all but three species (Table 2) was rejected because the nightly occurrence frequencies varied between 4 and 71%. Moreover, it was positively correlated with a taxon’s mean panel density (Fig. 2), indicating that low nightly occurrence was a result of the low overall taxon abundance. The low inter-species correlations in nightly densities (Fig. 3) indicate species-specific migration patterns rather than rare mass-migration events. Altogether, the results suggest that these taxa engaged in migration over many nights throughout the wet season, rather than during a few rare events, albeit certain nights, probably near peak activity, are of considerably higher density as is also reflected in the total insects caught (Fig. S3). The cross-panel occurrence (Table 2) indicates that clumping does not reflect tight flying “swarms.” Migration regularity was demonstrated by the similarity of the taxa composition (appearance) over multiple years (2013–2015) and in locations up to 100 km apart (Fig. 4 and Table 3). Thus, the analysis of taxa’s aerial density (Table 3) showed that 1) aerial density distribution was clustered even after accommodating variation due to season, year, village and altitude; and 2) the season (coarsely defined by 2–4 month periods), but not year or locality of sampling were statistically significant across taxa. Hence, the typical weather conditions of the wet season are suitable for these movements.

The magnitude of migration is illustrated as the number of insects expected to cross a 1km line perpendicular to the wind at altitude over a single night. Because our taxa were captured in altitudes spanning 40 to 290m agl, a conservative estimate of the depth of the flight layer is 200m. Using the average nightly wind speed (3.5m/s, Fig. 5c), we estimated the number of insects crossing this imaginary line throughout the night (14 hours sampling duration), yielding an average parcel of air of 176.4km length, 1km (width), and 0.2km (height). The average aerial density was calculated across all sampling nights (including zeros) during the migration season, which was estimated as the longest annual interval when migration occurred (days between the first and the last dates the taxon was captured, i.e., its “migration period.” The number of insects per taxa crossing over a single night 1km perpendicular to the wind 50 to 250m agl ranged between 7,800 (*Dysdercus* sp.) to 750,000 (*Ch. coletta*, Fig. 7). Extrapolating these values to the annual number of insects crossing the 1,000km line spanning Mali’s width at latitude 14.0°N suggests values between one hundred million (*Dysdercus* sp.) and 0.1 quadrillion (10^14^) (*Ch. coletta*). The mean total insect density/panel (280, Fig. S3) is >25 times greater than that of *Ch. coletta*, presently, our most abundant species (Table 2). These values dwarf the recently estimated number of insects flying above the United Kingdom (Hu et al., 2016). We believe that our estimates are conservative, but even if they overestimate the actual value by an order of magnitude, they underscore the enormous scale of these movements, and suggest that a sizeable fraction of the population engages in windborne migration.

**Fig. 7.**
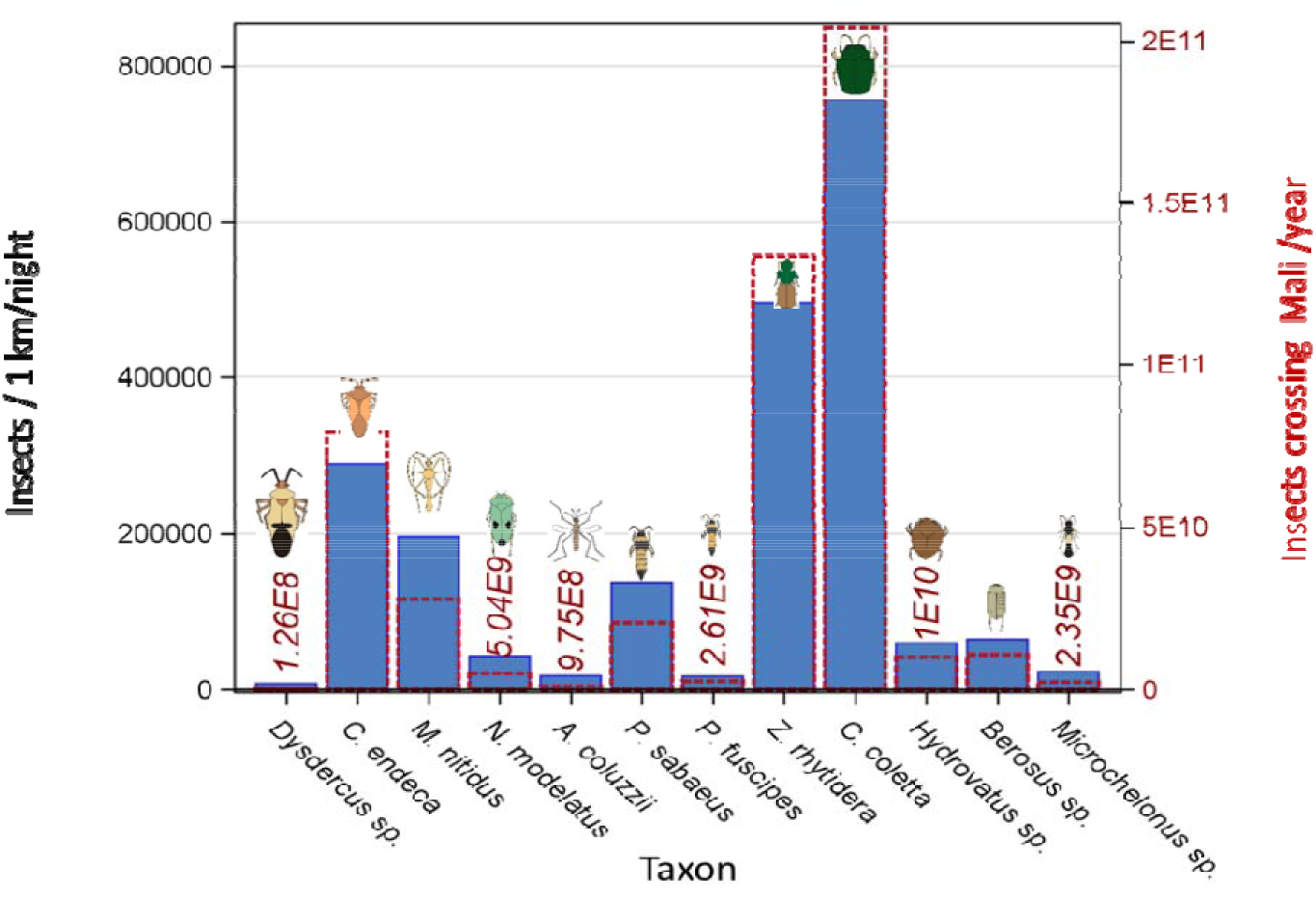
The number of insects per species crossing at altitude (50–250 m agl) imaginary lines perpendicular to the prevailing wind. Migrants per night per 1km (left Y axis, blue) are superimposed on the annual number per 1,000km line across Mali (right Y axis, red, see text).

Flight speeds of small insects (<3mg; Table S2) range around 1 m/s (Hocking, 1953; Snow, 1980), thus their overall displacement in altitude, where typical wind speed exceeds 4m/s (Fig. 5, S3, and below) is governed by the wind. The distance covered by windborne migrants depends on wind speed and the duration of their flights. Using a conservative average of wind speed at altitude of 4m/s (Fig. 5 and S3), an insect flying, for 2–10h would be transported 30– 140km, respectively (see below regarding entomological radar data showing small insects commonly flying in wind speeds >10m/s). Flight durations can be estimated by flight mills (Taylor et al., 2010; Lee & Leskey, 2015) or, less often, by the distance that they demonstrably flew and knowledge of the wind speed. Flight mill data suggest that leafhoppers (*N. virescens*) can fly over 10.75h (Cooter et al., 2000), similar to *A. gambiae* s.l. (10h) (Briegel et al., 2001; Kaufmann & Briegel, 2004).

Meteorological radar data from the Sahel reveal wind speed means between 8 and 12m/s, with occasional nights when wind speed in the lower jet stream (LJS, 150–500m agl) exceed 15m/s (Madougou et al., 2012). Reanalysis datasets such as MERRA-2 or ERA5 consistently underestimate wind speed in that layer (Fiedler et al., 2013). Because migrant insects often concentrate at the layer with maximal air speed (Drake & Reynolds, 2012), a small insect flying between 1 and 10 hours in a realistic average windspeed of the LJS, (10m/s) will cover on average 36–360km per night (over 500km in some nights). With trillions of insect migrants per season, many of which cover hundreds of kilometers in a single night - the implications of this behavior for ecosystem stability, public health, and especially for agriculture and food security are profound.

The seasonally productive habitats of the Sahel border diverse and “teeming” sub-equatorial habitats. This combination may result in a remarkable abundance of aerial insect migrants. This, in turn, may account for concentration of specialist aerial predators such as swifts (Åkesson et al., 2012, 2016), nightjars (Jackson, 2003), and bats (Fenton & Griffin, 1997) during peak migration period in this region. Recent reports on swifts reveal that during their fall migration, they arrive in the Sahel (latitudes 11–15°) in mid to late August when insect migration peaks, and that they remain in the area for ~ 24 d, 30% (9–67%) of their total migration duration, whilst covering only 9% of their total route. This contrasts with their spring migration in May, before insect high-altitude migration builds up, when they stay in the Sahel ~4 d, constituting only 14% (3–38%) of their journey (Åkesson et al., 2012). It appears that the swifts rely on the extreme insect abundance at altitude after crossing the Sahara and before heading to equatorial regions, where they overwinter. Their stations in-route support the remarkable magnitude of Sahelian insect migration, and suggest it is greater than that expected in equatorial regions and thus that it may represent a global hot zone for migratory insects.

The continuous seasonal movement of the Inter-Tropical Convergence Zone (ITCZ) implies shifting conditions and resources across the girth of the Sahel. During its short wet season a mosaic of patches receive high and low rainfall in any given year, which in turn reinforces migration (Southwood, 1962; Sellers, 1980; Drake & Gatehouse, 1995; Pedgley et al., 1995; Schneider et al., 2014). Movement between resource patches is predicted, especially for inhabitants of ephemeral water such as puddles, e.g., *A. coluzzii*. Marked seasonality with migration peaking during the rainy season when resources in the Sahel abound was evident in most taxa and might have been found in all taxa, had larger sample sizes been available (Table 3 and Fig. 3). Migration dynamics in nine of the thirteen taxa exhibited bimodal activity similar to the total density of insects/panel (Fig. S3), whereas only *A. coluzzii*, *Dysdercus* sp., and *Microchelonus* sp. exhibited a unimodal pattern (Fig. 3, Table 4). A wide unimodal migration peak fits the “residential Sahelian migration” strategy that promotes dispersal in species that persist in the Sahel throughout the year, but with migration into new “resource patches”. Such movement safeguards against severe wet-season droughts (Nicholson, 2013) that could eliminate local populations. The “residential Sahelian migration strategy” implies an ability to withstand the Sahelian dry season via aestivation (or quiescence) as is the case for *A. coluzzii* (Yaro et al., 2012; Dao et al., 2014; Huestis & Lehmann, 2014), although the mere existence of aestivation does not imply residential migration because aestivation may also occur in Savannas and Forest-Savanna zones (Masaki, 1980; Denlinger, 1986). Additionally, Oedaleus senegalensis (Acrididae) exhibits aestivation— eggs can survive several years in dry soil—but it can also cross the Sahel into the Savanna and return (over its 3 annual generations) (Cheke et al., 1990; Drake & Gatehouse, 1995; Maiga et al., 2008; Drake & Reynolds, 2012). Bi-modal flight activity better fits a “round-trip migration strategy”, whereby insects arrive in the Sahel from “perennial” habitats closer to the equator during the early peak, and “return” southwards when conditions in the Sahel are about to deteriorate with the approaching dry season, e.g., *Dysdercus volkeri* Schmidt in Ivory Coast and Mali (Duviard, 1977). Note that our *Dysdercus* sp. exhibited a uni-modal peak (Fig. 3), but the small number of specimens may have obscured a more complex pattern (Table 2). Presumably, one or more generations benefit from the lush conditions during the Sahelian wet season before embarking on their return flight, manifested by their second peak. The “round-trip migration” strategy predicts flight directions with a dominant northward component during the wet season and southward component as the dry season approaches. For example, *O. senegalensis* flies southward from the northern Sahel on strong Harmattan winds and covers 300–400 km in a night’s migration, although its northward movements seem gradual (Cheke et al., 1990; Pedgley et al., 1995; Drake & Reynolds, 2012). The late wet-season peak observed in October in total insect density/panel (Fig. S3) supports a rise in population density as predicted. However, contrary to prediction, wind direction during the seven nights with highest total insect density had a predominant northward component (not shown). These migration strategies are not mutually exclusive as species may exhibit both “round-trip migration” and “residential migration” in different populations. For example, equatorial populations of *A. coluzzii* (della Torre et al., 2002, 2005; Neafsey et al., 2015), are not expected to extensively engage in windborne migration. Possibly, species employing the “round trip” strategy may incorporate movements similar to “residential Sahelian migration” during the wet season, to better exploit the shifting rains and then return southwards to habitats with perennial resources, as exemplified by *O. senegalensis* which combines migration between the Sahel and the Savanna with a capacity for aestivation (above). Distinguishing among these possibilities and linking them to life history traits require additional information, which are currently are unverified for most Sahelian taxa (Table 4).

Wind directions during the period of flight activity spanned well over 180° for all taxa (Fig. 6 and Table 3), suggesting that movement between resource patches in the Sahel is widespread. During the rains, movements northwards and eastwards were especially common, following the ITCZ and its associated storms. Rains after the long dry season signifies high productivity, minimal competition, predation, and parasitism (Drake & Gatehouse, 1995). During the end of the wet season, movement southward was observed in 10 of the 12 taxa, however, there was no evidence of selective flight on southbound winds as was found over Europe (Chapman et al., 2010; Hu et al., 2016). The prevailing seasonal winds in the Sahel— humid southwest monsoon and dry northeast Harmattan—happen to take the migrants in seasonally appropriate directions, reducing the pressure for wind selectivity that has been demonstrated in temperate zones, where selectivity may confer much greater benefit. Additionally, important return migrations were possibly missed because we had fewer sampling nights during October–December (Fig. 1), and because during this time the LJS may have been higher than 190m and most insects flew above our traps.

In conclusion, our results demonstrate that a multitude of Sahelian insects regularly engage in high-altitude windborne migration in enormous densities throughout the rainy season. The dynamics of the studied taxa suggest species-specific drivers regardless of weather conditions during migration; however, the similarities among the taxa are striking. The dominant winds— southerly monsoon during the wet season and northerly Harmattan—during the dry season structure the general sources and destinations, yet, all taxa exploited winds that transported them to various directions and destinations, indicating an intra-Sahelian patch interchange. The specific drivers, mechanisms, and implications of these movements await to be revealed.

## Acknowledgements

We are grateful to the residents of Thierola, Siguima, Markabougou, and Dallowere for their permission to work near their homes, and for their wonderful assistance and hospitality. We thank to Dr. Moussa Keita, Mr. Boubacar Coulibaly, and Ousmane Kone for their valuable technical assistance with field and laboratory operations; Dr. Gary Fritz for his advice on the aerial sampling method; Drs. Thomas Henry, Warren Steiner, Saverio Rocchi, John Ascher, and Davide Dal Pos for their morphological identifications of *Dysdercus* sp. and *Cysteochila endeca*, *Hydrovatus* sp., *Hypotrigona* sp., and *Microchelonus* sp., respectively; and Dr. Terry Erwin for confirming the identification of *Zolotarevskyella rhytidera* by LC. We thank Julien Haran for his help with the identification of Smicronyxini weevils; Drs. Dick Sakai, Sekou F Traore, Jennifer Anderson, and Thomas Wellems, Ms. Margie Sullivan, and Mr. Samuel Moretz (National Institutes of Health, USA) for logistical support; Drs. Frank Collins and Neil Lobo (Notre Dame University, USA) for support to initiate the aerial sampling project; and Mr. Leland Graber for providing insect artwork used throughout the figures. We thank Drs. José MC Ribeiro and Alvaro Molina-Cruz (National Institutes of Health, USA) for reading earlier versions of this manuscript and providing us with helpful suggestions; and Drs. Alice Crawford and Fong (Fantine) Ngan, (NOAA/Air Resources Laboratory and CICS, the University of Maryland) for conversions of the MERRA2 to HYSPLIT format.

## Authors Contributions

This project was conceived by TL. Field methods and operations were designed by DLH with input from DRR and JWC. Fieldwork, protocol optimization, sampling data acquisition and management, and initial specimens processing was performed by AD, ASY, MD, DS, ZLS, and OY. Laboratory processing and identification of morphotypes was done by JF and LV with key inputs from LC, ET, HF, CM, MB, CB, and CS. Data entry and management was done by JF and LV. Morphological species identification and molecular analysis of specimens were conducted primarily by LC, ET, Y-ML, HF, CM, MB, CB, CS, RM, and RF. Data analysis were carried out by TL with inputs from all authors, especially BJK, RF, DRR, JWC, ES and Y-ML. The manuscript was drafted by TL JF and LV and revised by all authors. Throughout the project, all authors have contributed key ingredients and ideas that have shaped the work and the final paper.

## Data Deposition

The dataset will be deposited in D<u>ryad</u>.

---

Figure S1. Plates 1-18. 1: *Dysdercus* sp. (Pyrrhocoridae), 2, 3: *Cysteochila endeca* Drake (Tingidae), 4: *Metacanthus nitidus* Štusá (Berytidae), 5, 6: *Nephotettix modulatus* Melichar (Cicadellidae), 7: *Anopheles coluzzii* Coetzee & Wilkerson (male, Culicidae), 8: *Paederus sabaeus* Erichson, *Paederus fuscipes* Curtis (Staphylinidae), 9, 10: *Zolotarevskyella rhytidera* (Chaudoir) (Carabidae), 11, 12: *Chaetocnema coletta* Bechyn (Chrysomelidae), 13, 14: *Hydrovatus* sp. (Dytiscidae), 15, 16: *Berosus* sp. (Hydrophilidae), 17: *Microchelonus* sp. (Brachonidae), 18: *Hypotrigona* sp. (Apidae). Ruler units are mm, and a scale bar is shown on selected photos.

**Figure S2.**
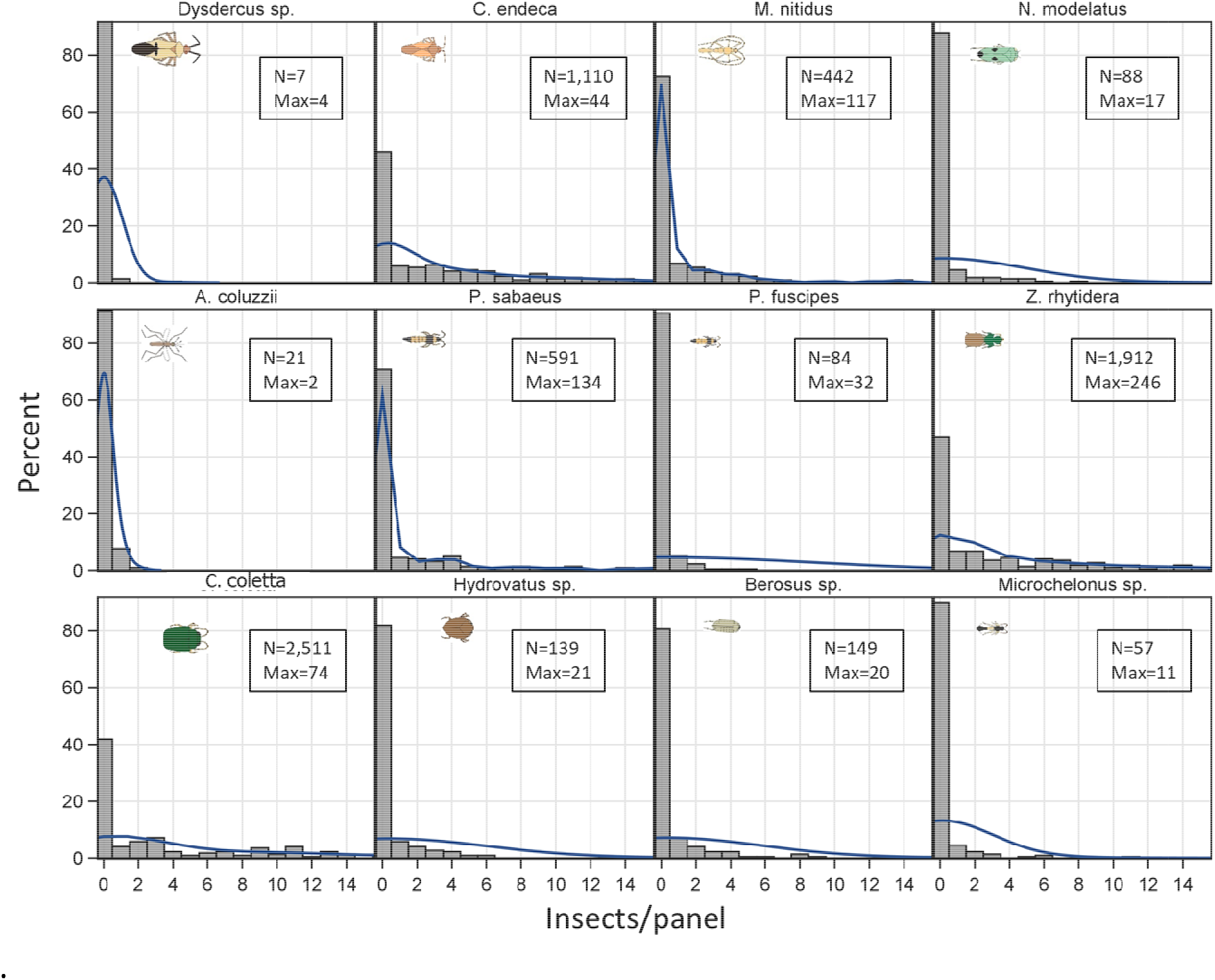
L-shape distributions of the number of insects per panel (histogram) and kernel density function (line) for the selected taxa. Maximum value of the X-axis was truncated to 15 (some values were higher, see Max value, inset). N and Max denote the total and the maximum density/panel in 220 sticky nets.

**Fig S3.**
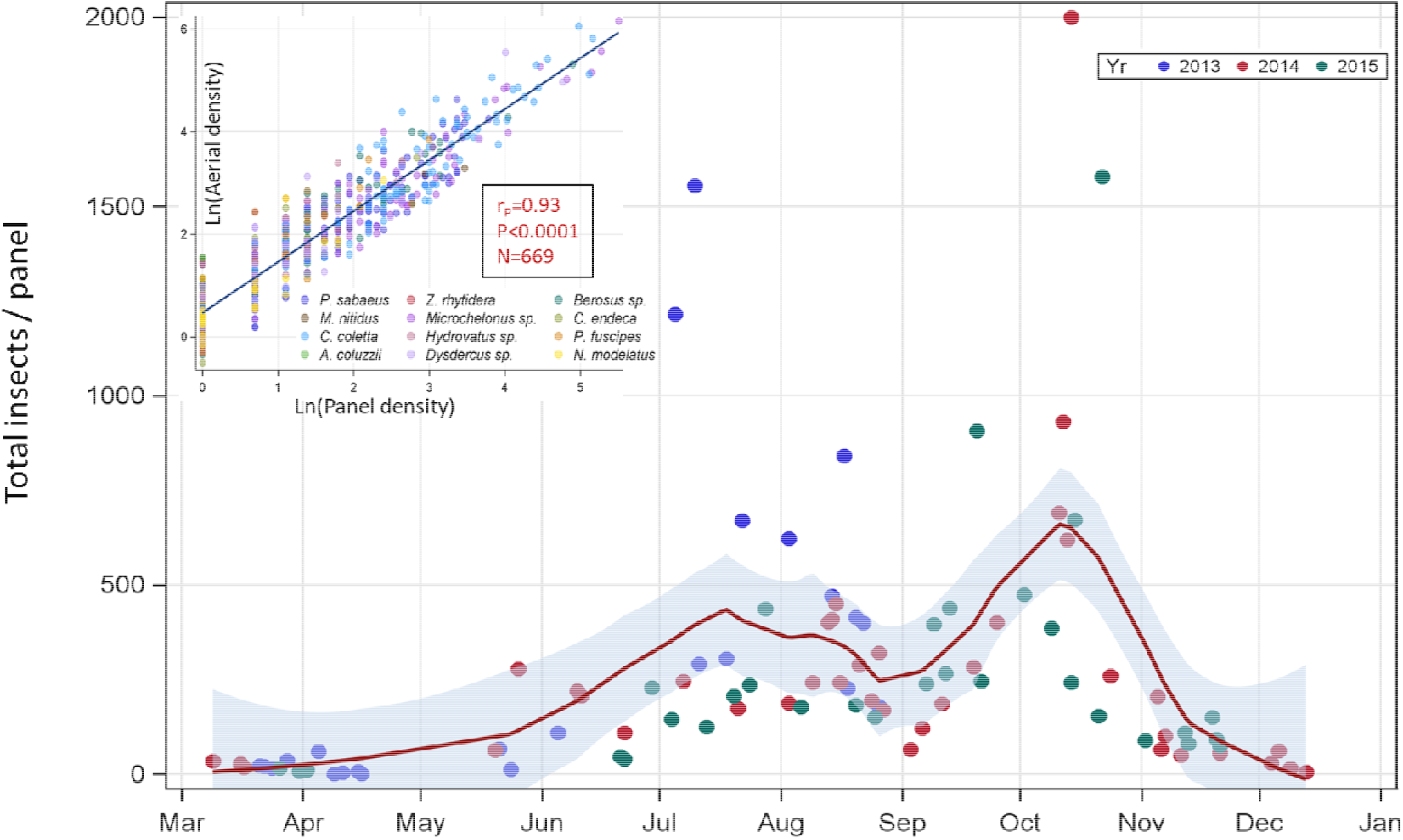
Seasonal variation in total-insect density/panel (note: sampling in 2013 ended in August). Inset: Relationship between panel density and aerial density (log scales). Pearson correlation coefficient (N reflects values larger than 0).

**Fig. S4.**
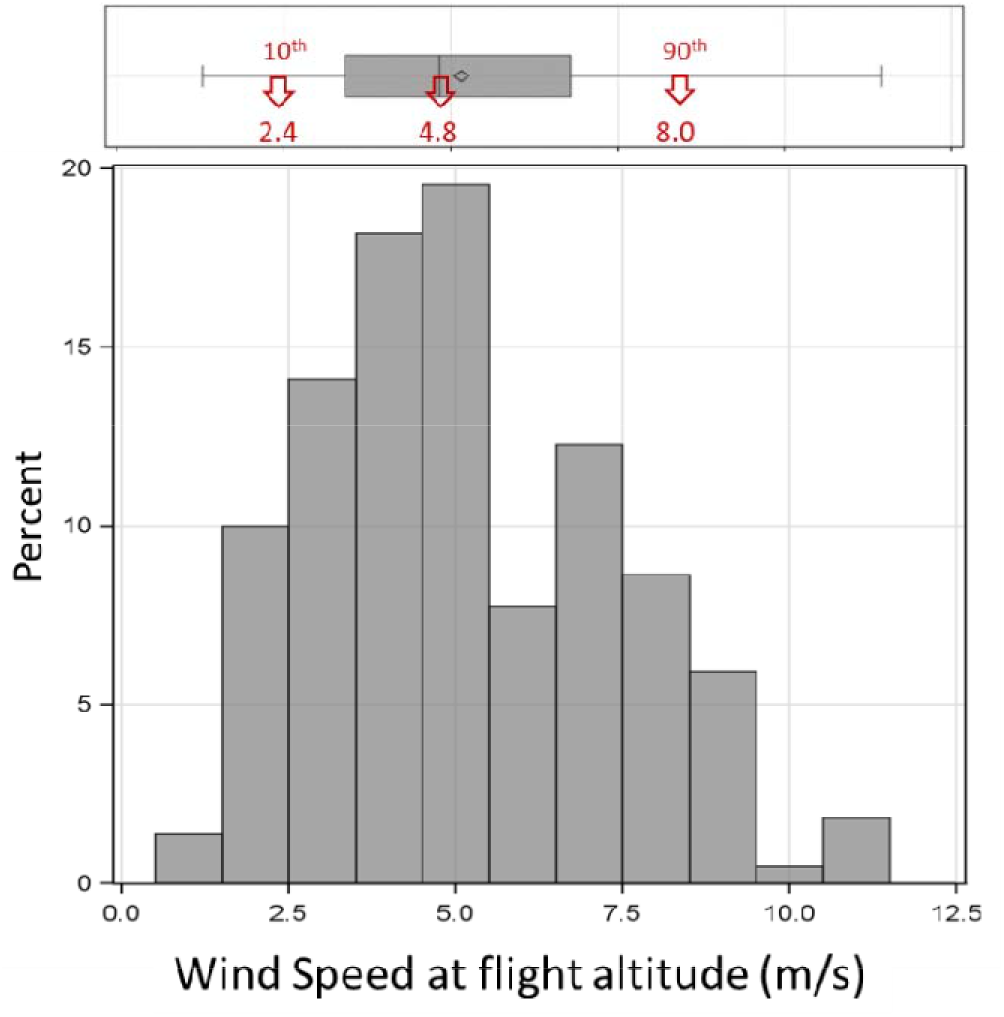
Distribution of mean nightly wind speed at flight height. Estimates are based on MERRA2 database and matched to the nearest panel altitude (50m, 70m and 200 m agl, see text from details).

