## Supplemental Fig. 1 for "Massive windborne migration of Sahelian insects: Diversity, seasonality, altitude, and direction"

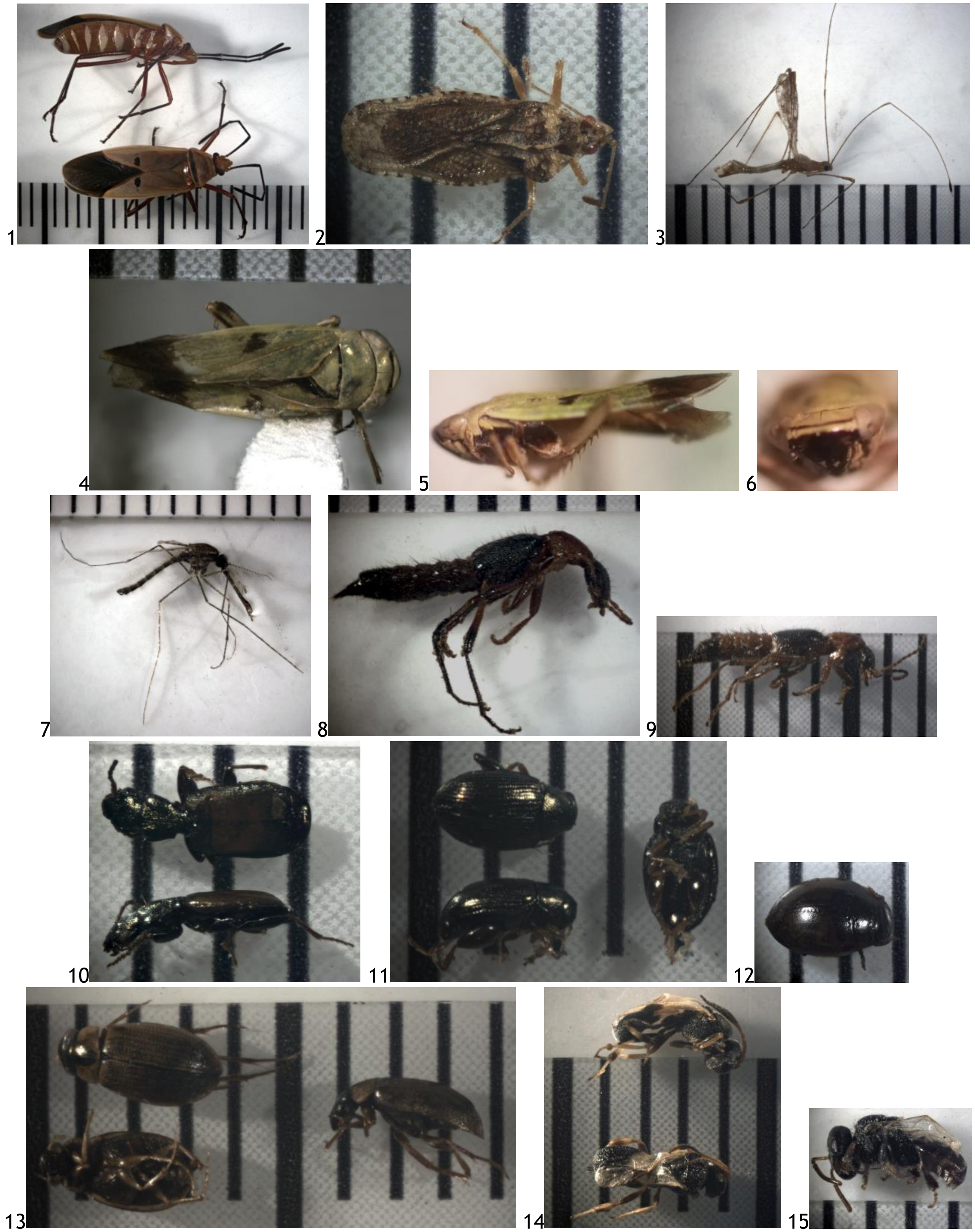

Plates 1-18. Selected species pictured with ruler; each line is 1mm. 1: *Dysdercus* sp., 2: *Cy. endeca*, 3: *M. nitidus*, 4-6: *N. modulatus*, 7: *A. coluzzii* (male), 8: *P. sabeus*, 9: *P. fuscipes*, 10: *Z. rhytidera*, 11: *Ch. coletta*, 12: *Hydrovatus* sp., 13: *Berosus* sp., 14: *Microchelonus* sp., 15: *Hypotrigona* sp.
