## Supplemental Table 1 for "Massive windborne migration of Sahelian insects: Diversity, seasonality, altitude, and direction"

| Collection Site ( | ollection Date | Height (m) | TubeIDOrig | Order | suborder | superfamily | family | subfamily | tribe | genus | species | TaxonomistName |
|---|---|---|---|---|---|---|---|---|---|---|---|---|
| Markabougou, Mali | 3-Aug-13 | 40 | MB231A | Coleoptera | Adephaga | Caraboidea | Carabidae | Cicindelinae | Cicindelini | Lophyra | senegalensis (Dejean, 1 | Schüle |
| Thierola, Mali (13.6 | 18-Aug-13 | 40 | TB386 | Coleoptera | Adephaga | Caraboidea | Carabidae | Harpalinae | Amblystomina | Amblystomus | dispar (Basilewski, 1951) | Schüle |
| Thierola, Mali (13.6 | 21-Mar-13 | 40 | TB204 | Coleoptera | Adephaga | Caraboidea | Carabidae | Harpalinae | Amblystomina | Amblystomus | katanganus (Burgeon, 1 | Schüle |
| Thierola, Mali (13.6 | 7-Sep-15 | 120 | TB960B | Coleoptera | Adephaga | Caraboidea | Carabidae | Harpalinae | Amblystomina | Amblystomus | latefasciatus (Basilewsky | Schüle |
| Suigima, Mali (14.1 | 21-May-13 | 40 | SB111A | Coleoptera | Adephaga | Caraboidea | Carabidae | Harpalinae | Amblystomina | Amblystomus | viridulus (Erichson, 1843 | Schüle |
| Suigima, Mali (14.1 | 21-May-13 | 40 | SB111A | Coleoptera | Adephaga | Caraboidea | Carabidae | Harpalinae | Chlaeniini | Callistochrous | baxi (Gory, 1833) | Schüle |
| Suigima, Mali (14.1 | 21-May-13 | 40 | SB111A | Coleoptera | Adephaga | Caraboidea | Carabidae | Harpalinae | Harpalinae | Afromizonus | tecospilus (Basilewsky, 1 | Schüle |
| Suigima, Mali (14.1 | 21-May-13 | 40 | SB111A | Coleoptera | Adephaga | Caraboidea | Carabidae | Harpalinae | Harpalini | Egadroma | discriminatis (Basilewsky | Schüle |
| Suigima, Mali (14.1 | 21-May-13 | 40 | SB111A | Coleoptera | Adephaga | Caraboidea | Carabidae | Harpalinae | Harpalini | Platymetopus | tesselatus (Basilewsky, 1 | Schüle |
| Thierola, Mali (13.6 | 7-Sep-15 | 120 | TB960A | Coleoptera | Adephaga | Caraboidea | Carabidae | Lebiinae | Lebiini | Metadromius | royi (Mateu, 1969) | Schüle |
| Thierola, Mali (13.6 | 21-Mar-13 | 40 | TB204 | Coleoptera | Adephaga | Caraboidea | Carabidae | Lebiinae | Lebiini | Singilis | bedimo (Anichtchenko, 2 | Schüle |
| Suigima, Mali (14.1 | 21-May-13 | 40 | SB111A | Coleoptera | Adephaga | Caraboidea | Carabidae | Lebiinae | Lebiini | Syntomus | submaculatus (Wollaston | Schüle |
| Thierola, Mali (13.6 | 14-Aug-14 | 160 | TB535B | Coleoptera | Adephaga | Caraboidea | Carabidae | Lebiinae | Odacanthini | Lasiocera | nitidula (Dejean, 1831) | Schüle |
| Thierola, Mali (13.6 | 7-Sep-15 | 120 | TB960A | Coleoptera | Adephaga | Caraboidea | Carabidae | Lebiinae | Pentagonicini | Pentagonica | elegans (Peringuey, 189 | Schüle |
| Thierola, Mali (13.6 | 7-Sep-15 | 120 | TB960A | Coleoptera | Adephaga | Caraboidea | Carabidae | Trechinae | Bembidiini | Polyderis | sp. | Schüle |
| Suigima, Mali (14.1 | 21-Nov-16 | 160 | SB574A | Coleoptera | Adephaga | Caraboidea | Carabidae | | | Sphaerotachys | tetraspilus (Solsky, 1874 | Schüle |
| Markabougou, Mali (13.9128, -6.3425) | | | MB231A | Coleoptera | Adephaga | Dytiscoidea | Dytiscidae | Hydroporinae | | Hydrovatus | | Saverio Rocchi |
| Markabougou, Mali | 20-Aug-13 | 40 | MB255A-CO-2 | Coleoptera | Adephaga | Dytiscoidea | Dytiscidae | Laccophilinae | Laccophilini | Laccophilus | | Warren Steiner |
| Markabougou, Mali | 13-Aug-14 | 190 | MB395A-CO-2 | Coleoptera | Adephaga | Dytiscoidea | Dytiscidae | | | | | Warren Steiner |
| Thierola, Mali (13.6 | 28-Oct-14 | 160 | TB682A-CO-1 | Coleoptera | Adephaga | Caraboidea | Carabidae | Brachininae | Brachinini | Brachinus | prob. dorsalis Dejean | Lourdes Chamorro |
| Thierola, Mali (13.6 | 4-Jul-15 | | TB820A | Coleoptera | Adephaga | Caraboidea | Carabidae | Lebiinae | Lebiini | Zolotarevskyella | prob. rhytidodera (Csiki) | Lourdes Chamorro |
| Thierola, Mali (13.6 | 7-Sep-15 | 120 | TB960A | Coleoptera | Adephaga | Caraboidea | Carabidae | Trechinae | Bembidiini | Tachyura | biplagiatus (Dejean, 1831 | Schüle |
| Thierola, Mali (13.6 | 7-Sep-15 | 120 | TB960A | Coleoptera | Adephaga | Caraboidea | Carabidae | Trechinae | Bembidiini | Tachyura | cfr. vagans (Peringuey, 1 | Schüle |
| Thierola, Mali (13.6 | 7-Sep-15 | 120 | TB960A | Coleoptera | Adephaga | Caraboidea | Carabidae | Trechinae | Bembidiini | Tachyura | fumicata (Motschulsky, 1 | Schüle |
| Thierola, Mali (13.6 | 18-Aug-13 | 40 | TB386 | Coleoptera | Adephaga | Caraboidea | Carabidae | Trechinae | Bembidiini | Tachyura | spec. | Schüle |
| Suigima, Mali (14.1 | 7-Jul-14 | 40 | SB286A-CO-8 | Coleoptera | Adephaga | Caraboidea | Carabidae | | | | | Leonid Friedman |
| Thierola, Mali (13.6583, -7.2155) | | | TB669A | Coleoptera | Polyphaga | Bostrichoidea | Bostrichidae | | | | | Laura Verú |
| Suigima, Mali (14.1 | 7-Jul-14 | 40 | SB286A-CO-8 | Coleoptera | Polyphaga | Chrysomeloidea | Brentidae | Galerucinae | | | | Leonid Friedman |
| Thierola, Mali (13.6583, -7.2155) | | | TB21A-CO-1 | Coleoptera | Polyphaga | Chrysomeloidea | Chrysomelidae | Bruchinae | Pachymerini | Caryedon? | | Laura Verú |
| Thierola, Mali (13.6 | 13-Aug-14 | 190 | TB533B-CO-5 | Coleoptera | Polyphaga | Chrysomeloidea | Chrysomelidae | Bruchinae | | | | Leonid Friedman |
| Markabougou, Mali | 15-Oct-14 | | MB446A-CO-1 | Coleoptera | Polyphaga | Chrysomeloidea | Chrysomelidae | Criocerinae | | | | Lourdes Chamorro |
| Markabougou, Mali | 20-Aug-13 | 40 | MB255A-CO-2 | Coleoptera | Polyphaga | Chrysomeloidea | Chrysomelidae | Eumolpinae | | | | Lourdes Chamorro |
| Markabougou, Mali | 7-Jul-14 | 120 | MB320A | Coleoptera | Polyphaga | Chrysomeloidea | Chrysomelidae | Galerucinae | Alticini | Aphthona | laevissima (Wollaston) | Maurizio Biondi |
| Markabougou, Mali | 7-Jul-14 | 120 | MB320A | Coleoptera | Polyphaga | Chrysomeloidea | Chrysomelidae | Galerucinae | Alticini | Aphthona | signatifrons (Wollaston) | Maurizio Biondi |
| Markabougou, Mali | 13-Aug-14 | 190 | MB395B-CO-3 | Coleoptera | Polyphaga | Chrysomeloidea | Chrysomelidae | Galerucinae | Alticini | Aphthona | sp. 1 | Furth |
| Markabougou, Mali | 7-Jul-14 | 120 | MB320A | Coleoptera | Polyphaga | Chrysomeloidea | Chrysomelidae | Galerucinae | Alticini | Aphthona | whitfieldi (Bryant) | Maurizio Biondi |
| Markabougou, Mali | 14-Aug-14 | 190 | MB398B | Coleoptera | Polyphaga | Chrysomeloidea | Chrysomelidae | Galerucinae | Alticini | Chaetocnema | coletta (Bechyné) | Maurizio Biondi |
| Markabougou, Mali | 24-Oct-14 | 120 | MB474A | Coleoptera | Polyphaga | Chrysomeloidea | Chrysomelidae | Galerucinae | Alticini | Longitarsus | sp. 1 | Maurizio Biondi |
| Markabougou, Mali | 7-Jul-14 | 120 | MB320A | Coleoptera | Polyphaga | Chrysomeloidea | Chrysomelidae | Galerucinae | Alticini | Longitarsus | sp. 2 | Maurizio Biondi |
| Suigima, Mali (14.1 | 10-Aug-14 | 190 | SB362A-CO-9 | Coleoptera | Polyphaga | Chrysomeloidea | Chrysomelidae | Galerucinae | Alticini | | | Maurizio Biondi |
| Markabougou, Mali | 9-Aug-14 | 190 | MB386A | Coleoptera | Polyphaga | Chrysomeloidea | Chrysomelidae | Galerucinae | Luperini | Afromaculepta | decemmaculata | Thomas Wagner |
| Thierola, Mali (13.6 | 16-Mar-14 | 40 | TB416A-CO-4 | Coleoptera | Polyphaga | Chrysomeloidea | Chrysomelidae | Galerucinae | Luperini | Monolepta | sp. 1 | Furth |
| Thierola, Mali (13.6583, -7.2155) | | | TB392A-CO-3 | Coleoptera | Polyphaga | Chrysomeloidea | Chrysomelidae | Galerucinae | Luperini | Monolepta | sp. 2 | Furth |
| Thierola, Mali (13.6 | 14-Oct-14 | 120 | TB639A | Coleoptera | Polyphaga | Chrysomeloidea | Chrysomelidae | Galerucinae | Luperini | Monolepta | sp. 3 | Furth |
| Thierola, Mali (13.6 | 5-Jul-13 | 40 | TB321A | Coleoptera | Polyphaga | Chrysomeloidea | Chrysomelidae | Galerucinae | | Panafrolepta | dahlmani | Thomas Wagner |
| Suigima, Mali (14.1 | 17-Aug-13 | 160 | SB179A | Coleoptera | Polyphaga | Cucujoidea | Coccinellidae | Coccidulini | | | | Shockley |
| Markabougou, Mali | 15-Oct-14 | | MB446A-CO-2 | Coleoptera | Polyphaga | Cucujoidea | Coccinellidae | | | | | |
| Markabougou, Mali | 13-Aug-14 | 190 | MB395A-CO-2 | Coleoptera | Polyphaga | Cucujoidea | Nitidulidae | | | | | Warren Steiner |
| Suigima, Mali (14.1 | 11-Sep-14 | 120 | SB444A-CO-1 | Coleoptera | Polyphaga | Cucujoidea | Phalacridae | | | | | Warren Steiner |
| Thierola, Mali (13.6 | 20-May-14 | | TB426B | Coleoptera | Polyphaga | Curculionoidea | Attelabidae | Rhynchitinae | | | sp. 1 | Lourdes Chamorro |
| Markabougou, Mali | 15-Aug-14 | 160 | MB397A-CO-9 | Coleoptera | Polyphaga | Curculionoidea | Brentidae | Nanophyinae | | Nanophyes | poss. errans | Lourdes Chamorro |
| Thierola, Mali (13.6 | 10-Aug-14 | 190 | TB527A-CO-5 | Coleoptera | Polyphaga | Curculionoidea | Brentidae | Nanophyinae | | Nanophyes | sp. 1 | Lourdes Chamorro |
| Thierola, Mali (13.6 | 18-Aug-13 | 40 | TB386A | Coleoptera | Polyphaga | Curculionoidea | Brentidae | Nanophyinae | | Nanophyes | sp. 2 | Lourdes Chamorro |
| Suigima, Mali (14.1 | 11-Sep-14 | 120 | SB444A-CO-6 | Coleoptera | Polyphaga | Curculionoidea | Brentidae | Apioninae | | | | Lourdes Chamorro |
| Suigima, Mali (14.1 | 7-Sep-15 | | SB725 | Coleoptera | Polyphaga | Curculionoidea | Curculionidae | Bagoinae | | Bagous | sp. 1 | Lourdes Chamorro |
| Thierola, Mali (13.6 | 16-Mar-14 | | TB418A | Coleoptera | Polyphaga | Curculionoidea | Curculionidae | Baridinae | | | sp. 1 | Lourdes Chamorro |
| Thierola, Mali (13.6 | 17-Mar-14 | 120 | TB420A | Coleoptera | Polyphaga | Curculionoidea | Curculionidae | Baridinae | Baridini | Baris | picturatus | Lourdes Chamorro |
| Suigima, Mali (14.1 | 21-Nov-16 | 160 | SB574A-CO-11 | Coleoptera | Polyphaga | Curculionoidea | Curculionidae | Baridinae | | | sp. 3 | Lourdes Chamorro |
| Markabougou, Mali | 22-Aug-13 | | MB262A | Coleoptera | Polyphaga | Curculionoidea | Curculionidae | Brachycerinae | | | sp. 1 | Lourdes Chamorro |
| Suigima, Mali (14.1 | 7-Jul-14 | | SB288A | Coleoptera | Polyphaga | Curculionoidea | Curculionidae | Brachycerinae | | | sp. 2 | Lourdes Chamorro |
| Thierola, Mali (13.6 | 7-Sep-15 | | TB960A | Coleoptera | Polyphaga | Curculionoidea | Curculionidae | Brachycerinae | | | sp. 3 | Lourdes Chamorro |
| Suigima, Mali (14.1 | 20-Jul-15 | 190 | SB662A | Coleoptera | Polyphaga | Curculionoidea | Curculionidae | Ceutorhynchinae | Ceutorhynchini | Nr. Neocoeliodes | sp. 1 | Lourdes Chamorro |

| | | | | | | | | | | | | |
|---|---|---|---|---|---|---|---|---|---|---|---|---|
| Thierola, Mali (13.6 | 13-Nov-15 | 120 | TB1095A-CO-17 | Coleoptera | Polyphaga | Curculionoidea | Curculionidae | Conoderinae | | *Lobotrachelus* | *sp. 1* | Lourdes Chamorro |
| Thierola, Mali (13.6 | 6-Sep-14 | 190 | TB593A-CO-19 | Coleoptera | Polyphaga | Curculionoidea | Curculionidae | Curculioninae | Ochyromerini | *Endaeus* | *sp. 1* | Lourdes Chamorro |
| Suigima, Mali (14.1 | 10-Jul-13 | 120 | SB127A-CO-5 | Coleoptera | Polyphaga | Curculionoidea | Curculionidae | Curculioninae | Rhamphini | | *sp. 1* | Lourdes Chamorro |
| Thierola, Mali (13.6 | 5-Jul-13 | 40 | TB321A | Coleoptera | Polyphaga | Curculionoidea | Curculionidae | Curculioninae | Smicronychini | *Afrosmicronyx* | *dorsomaculatus (Julien H* | Lourdes Chamorro |
| Markabougou, Mali | 26-Mar-15 | 190 | MB544A-CO-6 | Coleoptera | Polyphaga | Curculionoidea | Curculionidae | Curculioninae | Smicronychini | *Afrosmicronyx* | *umbrinus* | Lourdes Chamorro |
| Thierola, Mali (13.6 | 9-Mar-14 | 160 | TB397A | Coleoptera | Polyphaga | Curculionoidea | Curculionidae | Curculioninae | Smicronychini | *Sharpia* | *bella* | Lourdes Chamorro |
| Suigima, Mali (14.1 | 7-Jul-14 | 160 | SB288A | Coleoptera | Polyphaga | Curculionoidea | Curculionidae | Curculioninae | Smicronychini | *Smicronyx* | *gossypii (Haran)* | Lourdes Chamorro |
| Markabougou, Mali | 15-Aug-14 | 160 | MB397A-CO-8 | Coleoptera | Polyphaga | Curculionoidea | Curculionidae | Curculioninae | Smicronychini | *Smicronyx* | ***zambianus n. sp. (Haran*** | Julien Haran & Lo |
| Suigima, Mali (14.1 | 22-May-13 | 160 | SB113A-CO-7 | Coleoptera | Polyphaga | Curculionoidea | Curculionidae | Lixinae | Lixini | *Microlarinus* | *sp.* | Lourdes Chamorro |
| Suigima, Mali (14.1666, -7.2332) | | | SB573 | Coleoptera | Polyphaga | Curculionoidea | Curculionidae | Scolytinae | | | | Laura Verú |
| Markabougou, Mali | 13-Aug-14 | 190 | MB395A-CO-8 | Coleoptera | Polyphaga | Curculionoidea | Curculionidae | | | | | Lourdes Chamorro |
| Markabougou, Mali | 22-Aug-13 | | MB262A | Coleoptera | Polyphaga | Curculionoidea | Erorhinidae | Erirhininae | | | *sp. 1* | Lourdes Chamorro |
| Markabougou, Mali | 7-Jul-14 | | MB319A | Coleoptera | Polyphaga | Curculionoidea | Brentidae | Apioninae | | | *#1 female* | Lourdes Chamorro |
| Thierola, Mali (13.6 | 5-Jul-13 | | TB322C | Coleoptera | Polyphaga | Curculionoidea | Brentidae | Apioninae | | | *#1 male* | Lourdes Chamorro |
| Markabougou, Mali | 25-Sep-14 | | MB429E | Coleoptera | Polyphaga | Curculionoidea | Brentidae | Apioninae | | | *#2 female* | Lourdes Chamorro |
| Thierola, Mali (13.6 | 5-Jul-13 | | TB322A | Coleoptera | Polyphaga | Curculionoidea | Brentidae | Apioninae | | | *#2 male* | Lourdes Chamorro |
| Thierola, Mali (13.6 | 11-Oct-14 | | TB630A | Coleoptera | Polyphaga | Curculionoidea | Brentidae | Apioninae | | | *#3* | Lourdes Chamorro |
| Thierola, Mali (13.6 | 11-Oct-14 | | TB630A | Coleoptera | Polyphaga | Curculionoidea | Brentidae | Apioninae | | | *#4* | Lourdes Chamorro |
| Thierola, Mali (13.6 | 21-Jul-14 | | TB502B | Coleoptera | Polyphaga | Curculionoidea | Brentidae | Apioninae | | | *#5* | Lourdes Chamorro |
| Markabougou, Mali | 24-Oct-14 | | MB474A | Coleoptera | Polyphaga | Curculionoidea | Brentidae | Apioninae | | | *#6* | Lourdes Chamorro |
| Thierola, Mali (13.6 | 7-Jul-14 | | TB463D | Coleoptera | Polyphaga | Curculionoidea | Brentidae | Apioninae | | | *#7* | Lourdes Chamorro |
| Markabougou, Mali | 7-Jul-14 | | MB319A | Coleoptera | Polyphaga | Curculionoidea | Brentidae | Apioninae | | | *#8* | Lourdes Chamorro |
| Thierola, Mali (13.6 | 22-Oct-15 | | TB1056B | Coleoptera | Polyphaga | Curculionoidea | Brentidae | Apioninae | | *Conapium #1* | | Lourdes Chamorro |
| Thierola, Mali (13.6 | 22-Oct-15 | | TB1056B | Coleoptera | Polyphaga | Curculionoidea | Brentidae | Apioninae | | *Conapium #2* | | Lourdes Chamorro |
| Markabougou, Mali | 7-Jul-14 | | MB319A | Coleoptera | Polyphaga | Curculionoidea | Curculionidae | Curculioninae | | | *sp. 1* | Lourdes Chamorro |
| Thierola, Mali (13.6 | 7-Jul-14 | | TB463D | Coleoptera | Polyphaga | Curculionoidea | Brentidae | Apioninae | Piezotrachelini | | *#1* | Lourdes Chamorro |
| Markabougou, Mali | 26-Aug-13 | | MB272A | Coleoptera | Polyphaga | Curculionoidea | Brentidae | Apioninae | Pizeotrachelini | | *#2* | Lourdes Chamorro |
| Markabougou, Mali | 15-Aug-14 | 160 | MB397A-CO-1 | Coleoptera | Polyphaga | Elateroidea | Elateridae | | | | | Warren Steiner/Lo |
| Suigima, Mali (14.1 | 11-Sep-14 | 120 | SB444A-CO-1 | Coleoptera | Polyphaga | Hydrophiloidea | Hydrophilidae | Hydrophilinae | Berosini | *Berosus* | | Warren Steiner |
| Markabougou, Mali | 15-Aug-14 | 160 | MB397A-CO-1 | Coleoptera | Polyphaga | Hydrophiloidea | Hydrophilidae | | | | | Warren Steiner |
| Thierola, Mali (13.6 | 3-Aug-14 | | TB512A-CO-2 | Coleoptera | Polyphaga | Scarabaeoidea | Scarabaeidae | | | | | Lourdes Chamorro |
| Markabougou, Mali | 13-Aug-14 | 190 | MB395A-CO-2 | Coleoptera | Polyphaga | Staphylinoidea | Staphilinidae | Pselaphinae | | | | Warren Steiner |
| Suigima, Mali (14.1 | 10-Aug-14 | 190 | SB362A-CO-9 | Coleoptera | Polyphaga | Staphylinoidea | Staphylindae | | | | | Leonid Friedman |
| Suigima, Mali (14.1 | 19-Jul-14 | 90 | SB326A-CO-1 | Coleoptera | Polyphaga | Staphylinoidea | Staphylinidae | Aleocharinae | Lomechusini | *Diplopleurus* | | Dr. Jan Klimaszev |
| Thierola, Mali (13.6 | 13-Aug-14 | 160 | TB532A-CO-1 | Coleoptera | Polyphaga | Staphylinoidea | Staphylinidae | Aleocharinae | Lomechusini | *Paramyrmoecia* | bipustulata | Dr. Jan Klimaszev |
| Thierola, Mali (13.6 | 05-Aug-13 | 160 | TB374A-CO-3 | Coleoptera | Polyphaga | Staphylinoidea | Staphylinidae | Aleocharinae | Lomechusini | *Zyras (Ctenodonia)* | | Dr. Jan Klimaszev |
| Thierola, Mali (13.6 | 13-Aug-14 | 160 | TB532A-CO-1 | Coleoptera | Polyphaga | Staphylinoidea | Staphylinidae | Paederinae | Paederini | *Paederus* | *fuscipes (Curtis)* | Dr. Frank |
| Suigima, Mali (14.1 | 06-Sep-14 | 190 | SB431A-CO-1 | Coleoptera | Polyphaga | Staphylinoidea | Staphylinidae | Paederinae | Paederini | *Paederus* | *sabaeus (Erichson)* | Dr. Frank |
| Thierola, Mali (13.6 | 13-Aug-14 | 160 | TB532A-CO-1 | Coleoptera | Polyphaga | Staphylinoidea | Staphylinidae | Paederinae | Paederini | *Paederus* | | Dr.Frank |
| Thierola, Mali (13.6 | 13-Aug-14 | 160 | TB532A-CO-1 | Coleoptera | Polyphaga | Staphylinoidea | Staphylinidae | Staphylininae | Staphylinini | *Gabronthus* | *maritimus (Motschulsky)* | Dr. Frank |
| Thierola, Mali (13.6 | 14-Sep-14 | 160 | TB616A-CO-1 | Coleoptera | Polyphaga | Staphylinoidea | Staphylinidae | Staphylininae | Staphylinini | *Gabronthus* | | Dr. Frank |
| Markabougou, Mali | 10-Jul-13 | 120 | MB214A-CO-3 | Coleoptera | Polyphaga | Staphylinoidea | Staphylinidae | Staphylininae | Staphylinini | *Philonthus* | | Dr. Frank |
| Thierola, Mali (13.6 | 10-Aug-14 | 190 | TB527A-CO-6 | Coleoptera | Polyphaga | Tenebrionidae | Aderidae | | | | | Warren Steiner |
| Suigima, Mali (14.1 | 11-Sep-14 | 120 | SB444A-CO-7 | Coleoptera | Polyphaga | Tenebrionidae | Anthicidae | | | | | Warren Steiner |
| Markabougou, Mali (13.9128, -6.3425) | | | MB476B | Coleoptera | Polyphaga | Tenebrionoidea | Mordellidae | | | | | Laura Verú |
| Markabougou, Mali | 10-Jul-13 | 120 | MB214A-DI-2 | Diptera | Brachycera | Carnoidea | Chloropidae | | | *Epimadiza* | | Amnon Freidberg |
| Suigima, Mali (14.1 | 10-Aug-14 | 190 | SB362A-DI-1 | Diptera | Brachycera | Carnoidea | Chloropidae | | | | | John Ismay |
| Markabougou, Mali | 14-Aug-13 | 160 | MB242A-DI-3 | Diptera | Brachycera | Carnoidea | Milichiidae | Madizinae | | *Phyllomyza* | | Amnon Freidberg |
| Thierola, Mali (13.6583, -7.2155) | | | TB513B | Diptera | Brachycera | Diopsoidea | Diopsidae | Diopsinae | Diopsini | *Diopsis* | | |
| Markabougou, Mali | 14-Aug-14 | 190 | MB398A-DI-3 | Diptera | Brachycera | Empidoidea | Dolichopodidae | | | | | Igor Grichanov |
| Markabougou, Mali | 13-Aug-14 | 160 | MB394B-DI-3 | Diptera | Brachycera | Ephydroidea | Curtonotidae | | | *Curtonotum* | *saheliense Tsacas, 1977* | Ashley Kirk-Sprigg |
| Thierola, Mali (13.6 | 6-Sep-14 | 160 | TB593A-DI-5 | Diptera | Brachycera | Ephydroidea | Drosophilidae | Steganinae | Steganini | *Leucophenga* | | Amnon Freidberg |
| Thierola, Mali (13.6 | 13-Aug-14 | 190 | TB533A-DI-4 | Diptera | Brachycera | Ephydroidea | Drosophilidae | | | | | Shane McEvey |
| Markabougou, Mali | 7-Nov-14 | 160 | MB493A-DI-2 | Diptera | Brachycera | Ephydroidea | Ephydridae | Discomyzinae | Psilopini | *Psitola* | | Amnon Freidberg |
| Thierola, Mali (13.6 | 6-Sep-14 | 160 | TB593A-DI-4 | Diptera | Brachycera | Ephydroidea | Ephydridae | Hydrellinae | | *Notiphila* | | Amnon Freidberg |
| Markabougou, Mali | 10-Jul-13 | 120 | MB214A-DI-1 | Diptera | Brachycera | Ephydroidea | Ephydridae | | | | | Tadeusz Zatwarni |
| Markabougou, Mali | 19-Jul-14 | 190 | MB353A-DI-1 | Diptera | Brachycera | Lauxanioidea | Lauxaniidae | | | | | Stephen Gaimari |
| Thierola, Mali (13.6 | 13-Aug-14 | 190 | TB533B-DI-4 | Diptera | Brachycera | Muscoidea | Anthomyiidae | | | | | Verner Michelsen |
| Markabougou, Mali | 14-Aug-13 | 40 | MB240A-DI-4 | Diptera | Brachycera | Muscoidea | Muscidae | Muscinae | Muscini | *Musca* | | Amnon Freidberg |
| Markabougou, Mali | 14-Aug-13 | 40 | MB240A-DI-4 | Diptera | Brachycera | Muscoidea | Muscidae | Muscinae | Stomoxyini | | | Amnon Freidberg |
| Markabougou, Mali | 14-Aug-14 | 190 | MB398A-DI-2 | Diptera | Brachycera | Muscoidea | Muscidae | Phaoniinae | Atherigonini | *Atherigona* | | Burgert Muller |
| Markabougou, Mali | 14-Aug-13 | 40 | MB240A-DI-4 | Diptera | Brachycera | Muscoidea | Muscidae | | | | | Marcia Couri |
| Markabougou, Mali | 22-Aug-13 | 40 | MB261A-DI-1 | Diptera | Brachycera | Oestroidea | Calliphoridae | Chrysomyiinae | Rhiniini | *Rhynchomyia* | | Amnon Freidberg |

| | | | | | | | | | | | | |
|---|---|---|---|---|---|---|---|---|---|---|---|---|
| Markabougou, Mali | 14-Aug-13 | 40 | MB240A-DI-2 | Diptera | Brachycera | Oestroidea | Calliphoridae | Chrysomyiinae | Rhiniini | | | Amnon Freidberg |
| Markabougou, Mali | 14-Aug-13 | 160 | MB242A-DI-3 | Diptera | Brachycera | Oestroidea | Rhiniidae | | | | | Knut Rognes |
| Markabougou, Mali | 14-Aug-13 | 40 | MB240A-DI-4 | Diptera | Brachycera | Oestroidea | Tachinidae | | | | | Pierfilippo Cerretti |
| Suigima, Mali (14.1 | 19-Jul-14 | 90 | SB326A-DI-1 | Diptera | Brachycera | Platypezoidea | Phoridae | | | | | Amnon Freidberg |
| Markabougou, Mali | 10-Jul-13 | 120 | MB214A-DI-1 | Diptera | Brachycera | Sciomyzoidea | Sepsidae | | | | | Andrey Ozerov |
| Thierola, Mali (13.6 | 13-Aug-14 | 160 | TB532A-DI-1 | Diptera | Brachycera | Syrphoidea | Pipunculidae | | | | | Marc De Meyer |
| Thierola, Mali (13.6 | 13-Aug-14 | 190 | TB533B-DI-3 | Diptera | Brachycera | Tephritoidea | Lonchaeidae | Lonchaeinae | | *Silba* | | Amnon Freidberg |
| Thierola, Mali (13.6 | 24-Aug-14 | 190 | TB563B-DI-4 | Diptera | Brachycera | Tephritoidea | Platystomatidae | | | | | Andrew Whittingto |
| Thierola, Mali (13.6 | 6-Sep-14 | 160 | TB593A-DI-4 | Diptera | Brachycera | Tephritoidea | Tephritidae | Dacinae | Ceratidini | *Ceratitis* | | Marc De Meyer |
| Markabougou, Mali | 14-Aug-13 | 160 | MB242A-DI-4 | Diptera | Brachycera | Tephritoidea | Ulidiidae | Ulidiinae | Ulidiini | *Phsysiphora* | | Elena Kameneva |
| Thierola, Mali (13.6 | 13-Aug-14 | 160 | TB532B-DI-1 | Diptera | Nematocera | Chironomoidea | Ceratopogonidae | | | | | Amnon Freidberg |
| Markabougou, Mali | 14-Aug-13 | 160 | MB242A-DI-2 | Diptera | Nematocera | Chironomoidea | Chironomidae | | | | | Torbjørn Ekrem |
| Thierola, Mali (13.6 | 7-Jul-14 | 120 | TB461A | Diptera | Nematocera | Culicomorpha | Simuliidae | Simuliinae | | *Simulium* | *griseicolle* | Peter Adler |
| Thierola, Mali (13.6 | 6-Sep-14 | 190 | TB593A | Hemiptera | Auchenorrhyncha | Fulgoroidea | Delphacidae | Delphacinae | Delphacini | *Leptodelphax* | *maculigera* | Charles Bartlett |
| Thierola, Mali (13.6 | 17-Oct-14 | 190 | TB650A | Hemiptera | Auchenorrhyncha | Fulgoroidea | Delphacidae | Delphacinae | Delphacini | *Perkinsiella* | *dorsata* | Charles Bartlett |
| Thierola, Mali (13.6 | 9/14/2014 | 160 | TB616A | Hemiptera | Auchenorrhyncha | Fulgoroidea | Delphacidae | Delphacinae | Delphacini | *Sogatella* | *albofimbriata* | Charles Bartlett |
| Markabougou, Mali (13.9128, -6.3425) | | | MB395B-HO-5 | Hemiptera | Auchenorrhyncha | Fulgoroidea | Delphacidae | Delphacinae | Delphacini | *Sogatella* | *cf albofimbriata* | Charles Bartlett |
| Thierola, Mali (13.6 | 20-Aug-13 | 120 | TB393A-HO-2 | Hemiptera | Auchenorrhyncha | Fulgoroidea | Delphacidae | Delphacinae | Delphacini | *Sogatella* | *cf furcifera* | Charles Bartlett |
| Thierola, Mali (13.6 | 14-Sep-14 | 160 | TB616A | Hemiptera | Auchenorrhyncha | Fulgoroidea | Delphacidae | Delphacinae | Delphacini | *Sogatella* | *nigeriensis* | Charles Bartlett |
| Thierola, Mali (13.6583, -7.2155) | | | TB462C | Hemiptera | Auchenorrhyncha | Fulgoroidea | Delphacidae | Delphacinae | Delphacini | *Sogatella* | *vibex* | Charles Bartlett |
| Thierola, Mali (13.6583, -7.2155) | | | TB616A-HO-3 | Hemiptera | Auchenorrhyncha | Fulgoroidea | Delphacidae | Delphacinae | Delphacini | *Sogatella* | | Charles Bartlett |
| Thierola, Mali (13.6 | 13-Aug-14 | 190 | TB533A | Hemiptera | Auchenorrhyncha | Fulgoroidea | Delphacidae | Delphacinae | Delphacini | *Thriambus* | *strennus* | Charles Bartlett |
| Thierola, Mali (13.6 | 13-Aug-14 | 190 | TB533A | Hemiptera | Auchenorrhyncha | Fulgoroidea | Delphacidae | Delphacinae | Delphacini | *Toya* | *ceresensis* | Charles Bartlett |
| Thierola, Mali (13.6 | 14-Sep-14 | 160 | TB616A | Hemiptera | Auchenorrhyncha | Fulgoroidea | Delphacidae | Delphacinae | Delphacini | *Toya* | *tuberculosa* | Charles Bartlett |
| Thierola, Mali (13.6583, -7.2155) | | | TB527A-HO-1 | Hemiptera | Auchenorrhyncha | Fulgoroidea | Delphacidae | Delphacinae | Delphacini | | | Charles Bartlett |
| Thierola, Mali (13.6 | 6-Sep-14 | 160 | TB593A-HO-8 | Hemiptera | Auchenorrhyncha | Fulgoroidea | Delphacidae | Stenocraninae | | *Stenocranus* | | Tatina Novoselsky |
| Thierola, Mali (13.6583, -7.2155) | | | TB426A | Hemiptera | Auchenorrhyncha | Fulgoroidea | Flatidae | | | | | Charles Bartlett |
| Thierola, Mali (13.6583, -7.2155) | | | TB650A | Hemiptera | Auchenorrhyncha | Fulgoroidea | Ricaniidae | | | | | Charles Bartlett |
| Thierola, Mali (13.6 | 13-Aug-14 | 160 | TB532B | Hemiptera | Auchenorrhyncha | Membracoidea | Cicadellidae | Deltocephalinae | Chiasmini | *Exitianus* | *distanti* | Charles Bartlett |
| Thierola, Mali (13.6 | 3-Aug-14 | | TB512A-HO-1 | Hemiptera | Auchenorrhyncha | Membracoidea | Cicadellidae | Deltocephalinae | Chiasmini | *Exitianus* | | James Zahniser |
| Markabougou, Mali | 7-Nov-14 | 160 | MB493A | Hemiptera | Auchenorrhyncha | Membracoidea | Cicadellidae | Deltocephalinae | Chiasmini | *Nephotettix* | *modulatus* | Charles Bartlett |
| Thierola, Mali (13.6583, -7.2155) | | | TB752A-HO-4 | Hemiptera | Auchenorrhyncha | Membracoidea | Cicadellidae | Deltocephalinae | Chiasmini | *Nephotettix* | *cf modulatus* | Charles Bartlett |
| Thierola, Mali (13.6 | 13-Aug-14 | 190 | TB533A-HO-1 | Hemiptera | Auchenorrhyncha | Membracoidea | Cicadellidae | Deltocephalinae | Paralimnini | *Psammotettix* | | Tatina Novoselsky |
| Thierola, Mali (13.6 | 24-Aug-14 | 190 | TB563A | Hemiptera | Auchenorrhyncha | Membracoidea | Cicadellidae | Deltocephalinae | Paralimnini | | | James Zahniser |
| Thierola, Mali (13.6583, -7.2155) | | | TB650A | Hemiptera | Auchenorrhyncha | Membracoidea | Cicadellidae | Deltocephalinae | Vartini? | | | Charles Bartlett |
| Thierola, Mali (13.6583, -7.2155) | | | TB688A | Hemiptera | Auchenorrhyncha | Membracoidea | Cicadellidae | | | | | Charles Bartlett |
| Thierola, Mali (13.6 | 17-Oct-14 | 190 | TB650A | Hemiptera | Heteroptera | Cimicoidea | Nabidae | | | | | Carsten Morkel |
| Thierola, Mali (13.6 | 14-Oct-15 | 120 | TB1035A | Hemiptera | Heteroptera | Coreoidea | Rhopalidae | | | | | Carsten Morkel |
| Thierola, Mali (13.6 | 13-Aug-14 | 190 | TB533B-HE-1 | Hemiptera | Heteroptera | Coreoidea | Stenocephalidae | | | *Dicranocephalus* | | Thomas Henry |
| Thierola, Mali (13.6583, -7.2155) | | | TB533A-HO-4 | Hemiptera | Heteroptera | Corixoidea | Corixidae | | | | | Charles Bartlett |
| Thierola, Mali (13.6583, -7.2155) | | | TB534A | Hemiptera | Heteroptera | Gerroidea | Gerridae | Trepobatinae | | | | |
| Thierola, Mali (13.6 | 5-Nov-14 | 160 | TB688A | Hemiptera | Heteroptera | Gerroidea | Gerridae | | | | | Carsten Morkel |
| Thierola, Mali (13.6 | 19-Sep-14 | 120 | TB627A | Hemiptera | Heteroptera | Gerroidea | Veliidae | Microveliinae | Microveliini | *Microvelia* | *sp.* | Andreas Krüger |
| Thierola, Mali (13.6 | 13-Aug-14 | 160 | TB532A-PH-1 | Hemiptera | Heteroptera | Hydrometroidea | Hydrometridae | Hydromerinae | | *Hydrometra* | | Tatina Novoselsky |
| Thierola, Mali (13.6 | 10-Aug-14 | 190 | TB527A-HE-2 | Hemiptera | Heteroptera | Lygaeoidea | Berytidae | Metacanthinae | Metacanthini | *Metacanthus* | *nitidus* | Carsten Morkel |
| Thierola, Mali (13.6 | 24-Aug-14 | 190 | TB563A-HE-4 | Hemiptera | Heteroptera | Lygaeoidea | Berytidae | Metacanthinae | | *Yemma* | | Tatina Novoselsky |
| Thierola, Mali (13 | 19-Sep-14 | 190 | TB629A | Hemiptera | Heteroptera | Lygaeoidea | Geocoridae | Geocorinae | | *Geocoris* | *aff. erytrops* | Andreas Krüger |
| Markabougou, Mali | 19-Jul-14 | 190 | MB353A-HE-5 | Hemiptera | Heteroptera | Lygaeoidea | Lygaeidae | Orsillinae | Nysiini | *Nysius* | | Thomas Henry |
| Suigima, Mali (14.1 | 24-Oct-14 | 190 | SB488A-HE-3 | Hemiptera | Heteroptera | Lygaeoidea | Lygaeidae | Rhyparochromina | Drymini | *Stilbocoris* | | Tatina Novoselsky |
| Suigima, Mali (14.1 | 24-Oct-14 | 120 | SB486A-HE-2 | Hemiptera | Heteroptera | Lygaeoidea | Lygaeidae | | | | | Thomas Henry |
| Thierola, Mali (13.6 | 24-Aug-14 | 190 | TB563A-HE-4 | Hemiptera | Heteroptera | Lygaeoidea | Oxycarenidae | | | *Camptotelus* | | Tatina Novoselsky |
| Thierola, Mali (13.6 | 13-Aug-14 | 160 | TB532A-HE-1 | Hemiptera | Heteroptera | Lygaeoidea | Phyparochromida | Rhyparochromina | Rhyparochromini | *Beosus* | | Tatina Novoselsky |
| Thierola, Mali (13.6 | 14-Oct-15 | 120 | TB1035A | Hemiptera | Heteroptera | Lygaeoidea | Rhyparochromida | Rhyparochromina | Myodochini | *cf. Paromius* | | Carsten Morkel |
| Thierola, Mali (13.6 | 14-Oct-15 | 120 | TB1035A | Hemiptera | Heteroptera | Lygaeoidea | Rhyparochromida | Rhyparochromina | Myodochini | *Paromius* | *gracilis* | Carsten Morkel |
| Suigima, Mali (14.1 | 24-May-13 | 40 | SB120A | Hemiptera | Heteroptera | Lygaeoidea | Rhyparochromida | Rhyparochromina | Ozophorini | *Ethaltomarus* | *glabrosus (Scudder)* | Andreas Krüger/E |
| Markabougou, M | 21-Jul-14 | 190 | MB359A | Hemiptera | Heteroptera | Lygaeoidea | Rhyparochromida | Rhyparochromina | Stygnocorini | *cf.Lasiosomus* | *sp.* | Andreas Krüger |
| Suigima, Mali (14.1 | 19-Jul-14 | 120 | SB327A-HE-2 | Hemiptera | Heteroptera | Lygaeoidea | Rhyparochromidae | | | | | Thomas Henry |
| Thierola, Mali (13 | 19-Sep-14 | 190 | TB629A | Hemiptera | Heteroptera | Miroidea | Miridae | | | | | Andreas Krüger |
| Thierola, Mali (13 | 14-Aug-14 | 160 | TB535B | Hemiptera | Heteroptera | Notonectoidea | Notonectidae | Anisopinae | | *Anisops* | *spec.* | Andreas Krüger |
| Markabougou, Mali (13.9128, -6.3425) | | | MB261A-HO-4 | Hemiptera | Heteroptera | Notonectoidea | Notonectidae | | | | | Charles Bartlett |
| Suigima, Mali (14.1 | 24-Oct-14 | 190 | SB488A-HE-3 | Hemiptera | Heteroptera | Pentatomoidea | Cydnidae | Cydnidae | Geotomini | *Dallasiellus* | *discrepans* | Tatina Novoselsky |

| | | | | | | | | | | | | |
|---|---|---|---|---|---|---|---|---|---|---|---|---|
| Suigima, Mali (14.1 | 24-Oct-14 | 120 | SB486A-HE-1 | Hemiptera | Heteroptera | Pentatomoidea | Cydnidae | Cydninae | Geotomini | *Aethus* | | Thomas Henry |
| Suigima, Mali (14.1 | 14-Oct-15 | 120 | SB795A | Hemiptera | Heteroptera | Pentatomoidea | Cydnidae | | | | | Carsten Morkel |
| Suigima, Mali (14.1 | 24-Oct-14 | 120 | SB486A-HE-3 | Hemiptera | Heteroptera | Pentatomoidea | Pentatomidae | Pentatominae | Antestini | *Adria* | *parvula* | Thomas Henry |
| Suigima, Mali (14.1 | 14-Oct-15 | 120 | SB795A | Hemiptera | Heteroptera | Pentatomoidea | Pentatomidae | | | | | Carsten Morkel |
| Thierola, Mali (13.6 | 14-Jul-14 | 190 | TB484A | Hemiptera | Heteroptera | Pyrrhocoroidea | Pyrrhocoridae | Pyrrhocorinae | | *Dysdercus* | *sp.* | Andreas Krüger |
| Suigima, Mali (14.1 | 7-Jul-14 | 40 | SB286A-HE-5 | Hemiptera | Heteroptera | Reduvioidea | Reduviidae | | | | | Carsten Morkel |
| Thierola, Mali (13.6 | 14-Sep-14 | 160 | TB616A-HE-1 | Hemiptera | Heteroptera | Tingoidea | Tingidae | Tingidae | Tingini | *Cysteochila* | *endeca* | Thomas Henry |
| Suigima, Mali (14.1 | 7-Jul-14 | 40 | SB286A-HE-6 | Hemiptera | Heteroptera | Tingoidea | Tingidae | Tinginae | Tingini | *Dictyla* | | Tatina Novoselsky |
| Thierola, Mali (13.6 | 5-Nov-14 | 160 | TB688A | Hemiptera | Heteroptera | Tingoidea | Tingidae | | | | | Carsten Morkel |
| Markabougou, Mali (13.9128, -6.3425) | | | MB416B | Hemiptera | Sternorrhyncha | Aphidoidea | Aphididae | | | | | Laura Verú |
| Thierola, Mali (13.6583, -7.2155) | | | TB533A-HO-4 | Hemiptera | Sternorrhyncha | Psylloidea | | | | | | Charles Bartlett |
| Markabougou, Mali | 21-Aug-14 | 190 | MB416B | Hymenoptera | Apocrita | Apoidea | Apidae | Apinae | Meliponini | *Hypotrigona* | | Corey Smith/John |
| Thierola, Mali (13.6 | 3-Aug-14 | | TB512A-HY-2 | Hymenoptera | Apocrita | Apoidea | Crabronidae | | | | | Elijah Talamas |
| Thierola, Mali (13.6 | 24-Aug-14 | 190 | TB563B-HY-2 | Hymenoptera | Apocrita | Apoidea | Megachilidae | | | | | Elijah Talamas |
| Thierola, Mali (13.6 | 20-Aug-13 | 40 | TB392A-HY-2 | Hymenoptera | Apocrita | Apoidea | Sphecidae | | | | | Elijah Talamas |
| Markabougou, Mali | 15-Oct-14 | | MB446A-HY-1 | Hymenoptera | Apocrita | Apoidea | | | | | | Elijah Talamas |
| | | | | Hymenoptera | Apocrita | Chalcidoidea | Chalcicidae | Epitraninae | | *Epitranus* | | Bob Copeland |
| Thierola, Mali (13.6583, -7.2155) | | | TB562B | Hymenoptera | Apocrita | Chalcidoidea | Eulophidae | | | | | Jason Mottern |
| Markabougou, Mali | 20-Aug-13 | 40 | MB255A-HY-1 | Hymenoptera | Apocrita | Chalcidoidea | Eupelmidae | | | | | Elijah Talamas |
| Thierola, Mali (13.6583, -7.2155) | | | TB838A | Hymenoptera | Apocrita | Chalcidoidea | Eurytomidae | | | | | |
| Thierola, Mali (13.6 | 24-Aug-14 | 190 | TB563A-HY-3 | Hymenoptera | Apocrita | Chalcidoidea | | | | | | Elijah Talamas |
| Thierola, Mali (13.6 | 24-Aug-14 | 190 | TB563B-HY-3 | Hymenoptera | Apocrita | Chrysidoidea | Bethylidae | | | | | Elijah Talamas |
| Suigima, Mali (14.1666, -7.2332) | | | SB053 | Hymenoptera | Apocrita | Chrysidoidea | Chrysididae | | | | | Laura Verú |
| Thierola, Mali (13.6 | 22-Oct-15 | 120 | TB1056B | Hymenoptera | Apocrita | Chrysidoidea | Dryinidae | Gonatopodinae | | *Gonatopus* | | Massimo Olmi |
| Thierola, Mali (13.6583, -7.2155) | | | TB608A | Hymenoptera | Apocrita | Cynipoidea | Figitidae | Eucoilinae | | *Afrostilba* | | Jason Mottern |
| Markabougou, Mali | 14-Aug-14 | 160 | MB397A-HY-1 | Hymenoptera | Apocrita | Diaprioidea | Diapriidae | Diapriinae | Psilini | *Coptera* | | Elijah Talamas |
| Markabougou, Mali | 15-Aug-14 | 160 | MB397A-HY-1 | Hymenoptera | Apocrita | Diaprioidea | Diapriidae | | | | | Elijah Talamas |
| Markabougou, Mali | 19-Jul-14 | 160 | MB352A-HY-4 | Hymenoptera | Apocrita | Ichneumonoidea | Braconidae | Cheloninae | | | | Elijah Talamas |
| Markabougou, Mali | 19-Jul-14 | 160 | MB352A-HY-1 | Hymenoptera | Apocrita | Ichneumonoidea | Braconidae | | | | | Elijah Talamas |
| Thierola, Mali (13.6583, -7.2155) | | | TB608A | Hymenoptera | Apocrita | Ichneumonoidea | Ichneumonidae | Tersilochinae | | | | Robert Kula |
| Thierola, Mali (13.6583, -7.2155) | | | TB820A | Hymenoptera | Apocrita | Platygastroidea | Scelionidae | Scelioninae | Gryonini | *Gryon* | | Elijah Talamas |
| Thierola, Mali (13.6 | 10-Aug-14 | 190 | TB527A-HY-2 | Hymenoptera | Apocrita | Platygastroidea | Scelionidae | Scelioninae | | *Calliscelio* | | Elijah Talamas |
| Thierola, Mali (13.6 | 24-Aug-14 | 190 | TB563B-HY-3 | Hymenoptera | Apocrita | Platygastroidea | Scelionidae | Scelioninae | | *Dicroscelio* | | Elijah Talamas |
| Thierola, Mali (13.6 | 24-Aug-14 | 190 | TB563A-HY-2 | Hymenoptera | Apocrita | Platygastroidea | Scelionidae | Scelioninae | | *Fusicornia* | *eos* | Elijah Talamas |
| Thierola, Mali (13.6 | 24-Aug-14 | 190 | TB563B-HY-3 | Hymenoptera | Apocrita | Platygastroidea | Scelionidae | Teleasinae | Teleasini | *Trimorus* | | Elijah Talamas |
| Markabougou, Mali | 22-Aug-13 | 40 | MB261A-HY-1 | Hymenoptera | Apocrita | Vespoidea | Formicidae | Myrmicinae | Crematogastrini | *Crematogaster* | | Brendon Boudinot |
| Suigima, Mali (14.1 | 2-Sep-14 | 190 | SB419A-HY-1 | Hymenoptera | Apocrita | Vespoidea | Formicidae | Myrmicinae | Crematogastrini | *Tetramorium* | | Brendon Boudinot |
| Thierola, Mali (13.6 | 3-Aug-14 | | TB512A-HY-2 | Hymenoptera | Apocrita | Vespoidea | Formicidae | Myrmicinae | Stenammini | *Messor* | | Brendon Boudinot |
| Markabougou, Mali | 14-Aug-13 | 160 | MB242A-HY-3 | Hymenoptera | Apocrita | Vespoidea | Formicidae | Ponerinae | Ponerini | *Anochetus* | | Brendon Boudinot |
| Thierola, Mali (13.6 | 28-Oct-14 | 160 | TB682A-HY-1 | Hymenoptera | Apocrita | Vespoidea | Formicidae | Ponerinae | Ponerini | *Brachyponera* | | Brendon Boudinot |
| Suigima, Mali (14.1 14-Aug-13 | | 160 | SB173A-HY-1 | Hymenoptera | Apocrita | Vespoidea | Formicidae | | | | | |
| Thierola, Mali (13.6 | 5-Aug-13 | 40 | TB372A-HY-2 | Hymenoptera | Apocrita | Vespoidea | Pompilidae | | | | | |
| Thierola, Mali (13.6 | 3-Aug-14 | | TB512A-HY-2 | Hymenoptera | Apocrita | Vespoidea | Rhopalosomatidae | | | | | Elijah Talamas |
| Thierola, Mali (13.6 | 2-Nov-15 | 120 | TB1062A | Neuroptera | Hemerobiiformia | Chrysopoidea | Chrysopidae | Chrysopinae | Chrysopini | *Brinckochrysa* | | Stephen J Brooks |
| Thierola, Mali (13.6 | 7-Jul-14 | 160 | TB462B | Neuroptera | Hemerobiiformia | Chrysopoidea | Chrysopidae | Chrysopinae | Chrysopini | *Chrysoperla* | *congrua* | Stephen J Brooks |
| Suigima, Mali (14.1666, -7.2332) | | | SB113A | Neuroptera | Hemerobiiformia | Mantispoidea | Mantispidae | | | | | Laura Verú |
| Thierola, Mali (13.6 | 22-Jul-13 | 120 | TB361A-OR-1 | Orthoptera | Caelifera | Acridoidea | Acrididae | Gomphocerinae | | | | Hojun Song |
| Thierola, Mali (13.6 | 22-Jul-13 | 120 | TB361A-OR-1 | Orthoptera | Caelifera | Acridoidea | Acrididae | Oedipodinae | | | | Hojun Song |
| Thierola, Mali (13.6 | 5-Aug-13 | 40 | TB372A-OR-2 | Orthoptera | Caelifera | Acridoidea | Acrididae | | | | | Hojun Song |
| Thierola, Mali (13.6 | 23-Jul-13 | 120 | TB364A-OR-1 | Orthoptera | Caelifera | Pyrgomorphoidea | Pyrgomorphidae | Pyrgomorphinae | Atractomorphini | *Atractomorpha* | | Ricardo Marino-Pe |
| Thierola, Mali (13.6 | 19-Jul-13 | 40 | TB351A-OR-1 | Orthoptera | Caelifera | Pyrgomorphoidea | Pyrgomorphidae | Pyrgomorphinae | Pyrgomorphini | *Pyrgomorpha* | | Ricardo Marino-Pe |
| Thierola, Mali (13.6 | 13-Oct-15 | 190 | TB1033A-OR-2 | Orthoptera | Caelifera | Tetrigoidea | Tetrigidae | Tetriginae | Tetrigini | | | Hojun Song |
| Thierola, Mali (13.6 | 28-Oct-14 | 190 | TB683A-OR-2 | Orthoptera | Ensifera | Grylloidea | Gryllidae | Oecanthinae | Oecanthini | *Oecanthus* | | Song Lab |
| Thierola, Mali (13.6 | 13-Oct-15 | 190 | TB1033A-OR-1 | Orthoptera | Ensifera | Tettigonioidea | Tettigoniidae | Conocephalinae | | | | Derek A. Woller & |
| Thierola, Mali (13.6 | 12-Oct-14 | 120 | TB633A-OR-1 | Orthoptera | Ensifera | Tettigonioidea | Tettigoniidae | Phaneropterinae | | | | Derek A. Woller & |
| Markabougou, Mali | 7-Nov-14 | 160 | MB493A-OR-1 | Orthoptera | Ensifera | Tettigonioidea | Trigonidiidae | Trigonidiinae | Trigonidiini | | | Hojun Song |
| Thierola, Mali (13.6583, -7.2155) | | | TB349I | Neuroptera | Myrmeleontiformi | Myrmeleontoidea | Myrmeleontidae | | | | | Laura Verú |
| Thierola, Mali (13.6583, -7.2155) | | | TB502A | Thysanoptera | | | | | | | | Laura Verú |
| Thierola, Mali (13.6583, -7.2155) | | | | | | | Culicidae | | | | | |
